# Ev-OSMOSE: An eco-genetic marine ecosystem model

**DOI:** 10.1101/2023.02.08.527669

**Authors:** Alaia Morell, Yunne-Jai Shin, Nicolas Barrier, Morgane Travers-Trolet, Bruno Ernande

## Abstract

In the last decade, marine ecosystem models have been increasingly used to project interspecific biodiversity under various global change and management scenarios, considering ecological dynamics only. However, fish populations may also adapt to climate and fishing pressures, via evolutionary changes, leading to modifications in their life-history that could either mitigate or worsen, or even make irreversible, the impacts of these pressures. Building on the multispecies individual-based model Bioen-OSMOSE, an eco-evolutionary fish community model, Ev-Osmose, has been developed to account for evolutionary dynamics together with physiological and ecological dynamics in fish diversity projections. A gametic inheritance module describing the individuals’ genetic structure has been implemented. The genetic structure is defined by finite numbers of loci and alleles per locus that determine the genetic variability of growth, maturation and reproductive effort. Climate change and fishing activities will generate selection pressures on fish life-history traits that will respond through microevolution. This paper is an overview of the Ev-OSMOSE model. To illustrate the ability of the Ev-OSMOSE model to represent realistic fish community dynamics, genotypic and phenotypic traits’ mean and variance and consistent evolutionary patterns, we applied the model to the North Sea ecosystem. The simulated outputs are confronted to observed data of commercial catch, maturity ogives and length at age and to estimates of biomass for each modeled species. In addition to the evaluation of their mean value, the emerging traits’ variability is confronted to length-at-age and maturity data. To ensure the consistency of genetic inheritance and the resulting evolutionary patterns, we assessed the transmission of traits’ genotypic value across cohorts. Overall, the state of the modelled ecosystem was convincing at all these different biological levels. These results open perspectives for using Ev-OSMOSE in different marine regions to project the eco-evolutionary impact of various global change and management scenarios on different biological levels.

## 1. Introduction

Anthropogenic activities alter the ecological and evolutionary dynamics of marine ecosystems. In addition to inducing direct mortality, selective pressures such as fishing exploitation and climate change trigger changes in the life history traits of marine organisms due to evolutionary processes (Crozier and Hutchings, 2014; Heino et al., 2015). Knowledge about genetic diversity, its erosion, and its impact on organisms’ traits has been identified as a gap in current knowledge (IPBES, 2019), while existing studies have shown that small changes in traits, which may be evolutionary in nature, can imply large demographic and whole-community and ecosystem changes, with potential consequences for human activities (Audzijonyte et al., 2014, 2013). Incorporating genetic diversity and the resulting potential for adaptation into marine ecosystem models (MEMs) is thus considered as a key future development (Heymans et al., 2020; Rose et al., 2010). New modelling frameworks are needed to properly account for evolutionary changes and their impacts at the ecosystem scale to improve the reliability of predictions (Naish and Hard, 2008).

The existing marine modeling studies for addressing human-induced evolution have primarily focused on fisheries-induced evolution. The main modeling framework in this field is the eco-genetic model (Dunlop et al., 2009; Heino et al., 2015). Eco-genetic models are single species models that describe the individual’s life history, genetic variability using a quantitative genetic approach, density dependence and fishing as a selection pressure. More generally, this modelling framework allows the study of any pressure that induces evolutionary changes in life history traits (e.g, climate change, see Waples and Audzijonyte, 2016). However, eco-genetic models generally apply to single species, rendering difficult the upscale to the community and ecosystem levels, for example by accounting for the multiple interspecies interactions and the potential selective pressures those interactions may induce.

OSMOSE is a spatially explicit, multi-species and individual-based modeling framework for regional marine ecosystems (Shin and Cury, 2004). It includes the marine high trophic level components (fish and macro invertebrate) and fishing pressure explicitly and it is forced by coupled physical-biogeochemical models to represent the entire ecosystem. In this paper, we describe Ev-OSMOSE, a new modelling framework that incorporates an eco-genetic sub-model into OSMOSE. The eco-genetic sub-model explicitly describes individual genetic and phenotypic variability in life history traits for multiple species interacting in a food web. A bio-energetic sub-model describing the life history in response to biotic and abiotic conditions has already been integrated in the OSMOSE model resulting in a multispecies framework with a mechanistic modeling of life history (Bioen-OSMOSE model, Morell et al., 2023). Our new framework Ev-OSMOSE includes the genetic and phenotypic variances of life history traits described by the bio-energetic sub-model and thus allows the description of life history micro-evolution and adaptation in response to pressures. To our knowledge, this new model is the first marine ecosystem model to take into account micro-evolution and adaptation. This framework allows the study of evolutionary and ecological dynamics and their interactions at the multi-species level. It also allows to address the impacts of predation, fishing and climate-induced evolution. Featuring genetic variability, life history evolution, and multispecies interactions in a single framework make the model suitable for projecting future genetic, functional, and species diversity under fisheries and climate change scenarios, with consistent mechanisms linking these three organizational levels of biodiversity.

In this paper, we provide a detailed description of the principles and equations of the Ev-OSMOSE framework. Parameterization guidelines are provided with an application to the North Sea ecosystem as a case study example. Results from the North Sea application are also provided to verify the consistency of the new model developments.

## 2. Materials and methods

### 2.1. Model description

The Ev-OSMOSE model represents the eco-evolutionary dynamics of fish communities in marine ecosystems (Fig. 1). It is an individual-based, spatially-explicit multispecies model accounting for trophic interactions. The main characteristics of the model are opportunistic predation based on length and spatial co-occurrence of predators and prey, the mechanistic description of individuals’ life-history traits emerging from genetics and bioenergetics and the consideration of inter-individual phenotypic variability due to both genotypic variability and plastic responses to spatiotemporal variations in biotic and abiotic factors. The aim of the model is to explore the functioning and the eco-evolutionary dynamics of marine trophic webs, notably in response to perturbations such as fishing or climate change. The consequences of perturbations can be tracked from the individual genotype to the phenotype, to the population and to the community scale. The Ev-OSMOSE model extends the existing OSMOSE model by (i) explicitly accounting for the dependence of life-history traits on bioenergetics that, in turn, are determined by individual’s genotype, (ii) describing intra- and inter-specific genetic and abiotic phenotypic variability.

**Figure 1:**
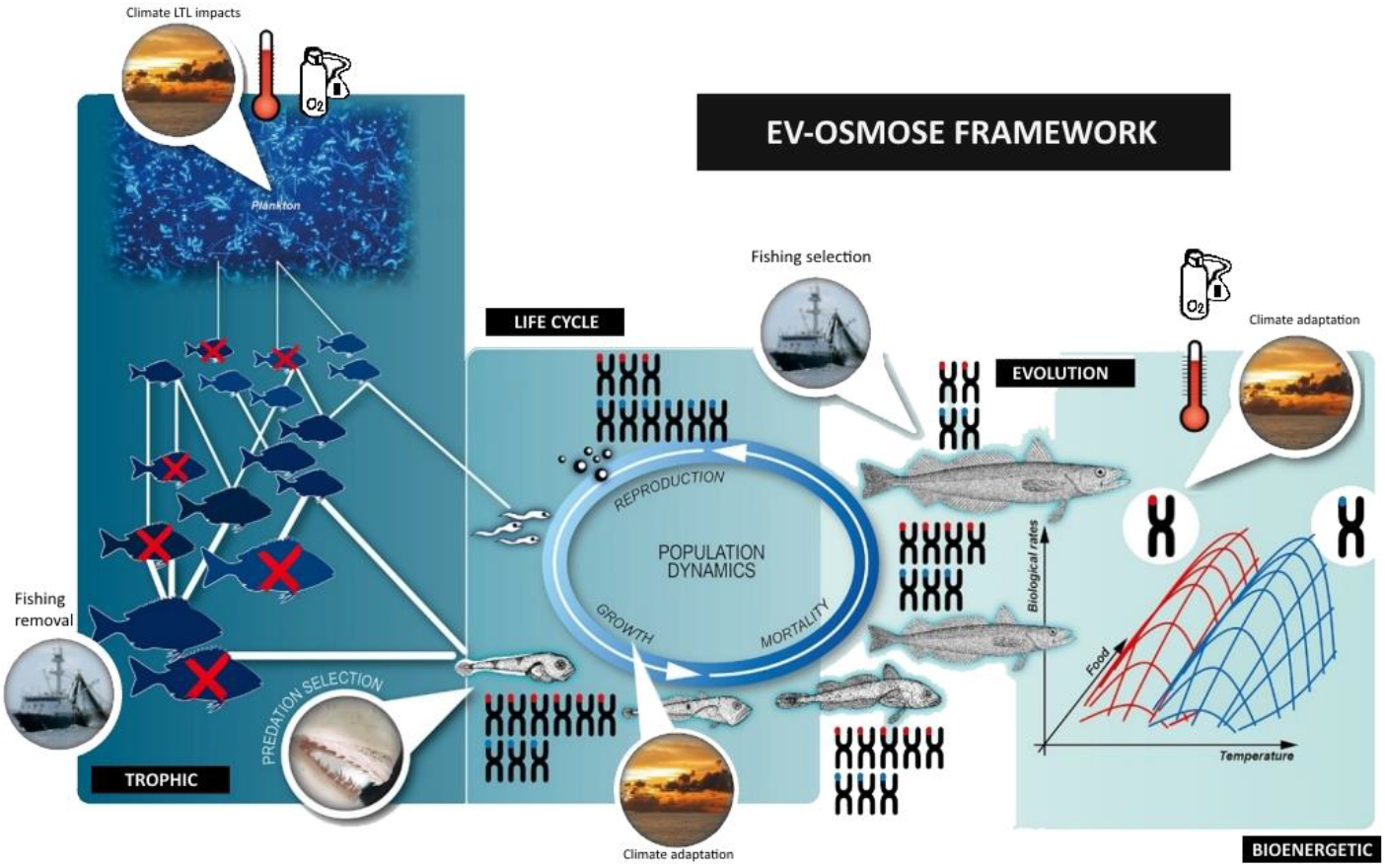
Graphical summary of the Ev-OSMOSE model. The Ev-OSMOSE model is a marine trophic web model where the trophic relationships emerge from species distributions per ontogenetic stage, spatiotemporal prey-predator co-occurrence and lengths adequacy, low trophic level (phytoplankton and zooplankton) biomass and species life cycle which is genetically determined and varies with temperature and oxygen.

**Figure 1:**
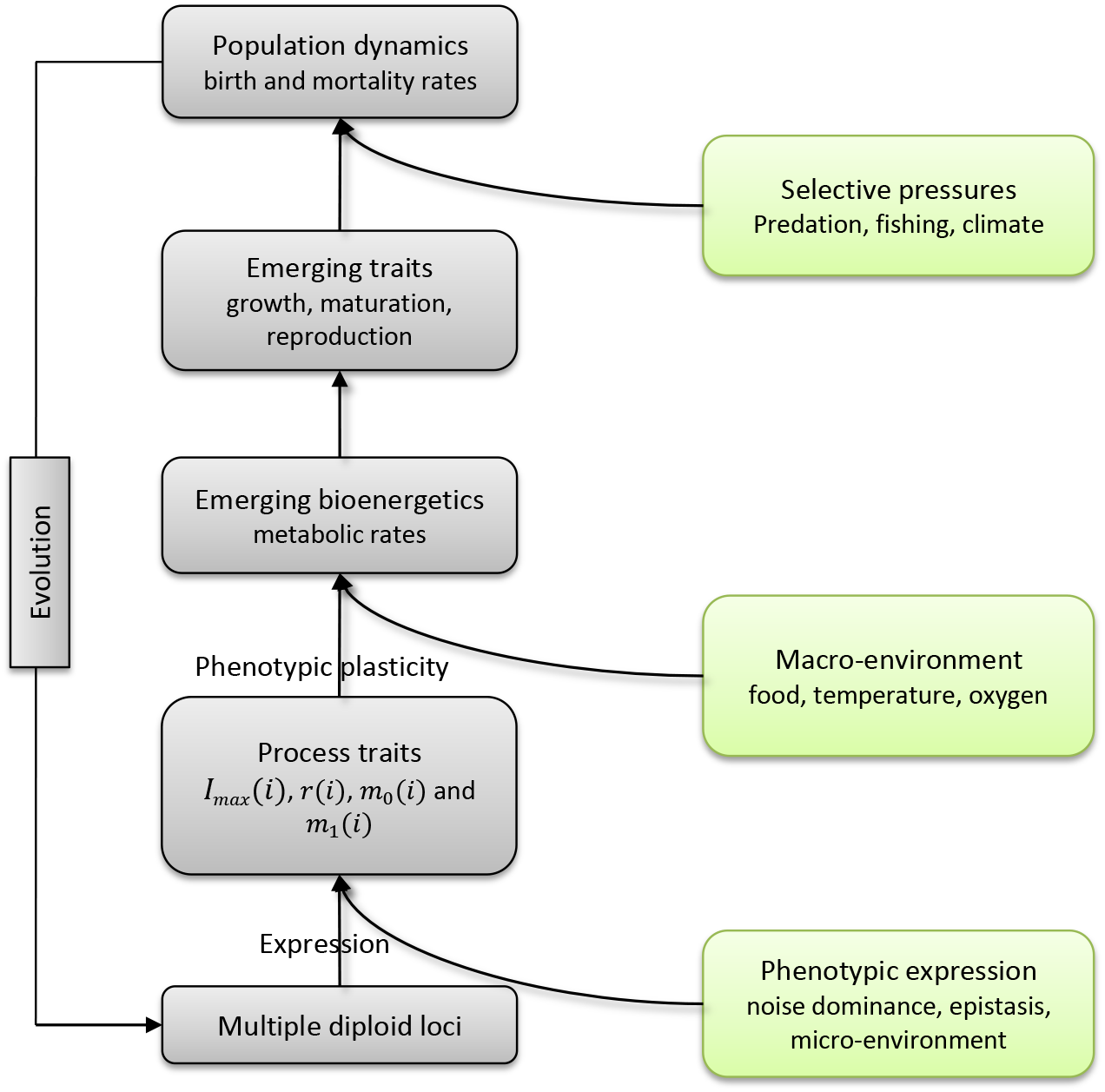
Ev-OSMOSE processes describing trait variations from loci to population level (grey) and the causes impacting trait values (green). The process traits are the traits underlying the physiological processes and trade-offs.

#### 2.1.1. Biological unit, state variables and spatial characteristics

The biological unit of the model is a school (a super-individual in individual-based modeling terms). It is formed of individuals from the same species that are biologically identical, i.e., whose state variables have the same values. Individuals are all diploid hermaphrodites, i.e. males and females are not distinguished, although the model is based on female life history. The state variables characterizing a school *i* at time step *t* belong to five categories (see Table 1 for parameters’ definitions and units and Table 2 for variables’ definitions and units):

- Trait genetic determinism and expression that include individuals’ genotype, composed of 2 alleles *A*_*z,l*,1_(*i*) and *A*_*z,l*,2_ (*i*) at each of the *l_z_* functional locus coding for each evolving trait *z* and 2 alleles *b*_*l*,1_(*i*) and *b*_*l*,2_(*i*) at each of *l_b_* neutral locus, and the phenotypic expression noise *e_z_*(*i*) for each evolving trait *z*;
- Ontogenic state of individuals described by their age *a*(*i, t*), somatic mass *w*(*i,t*) and gonadic mass *g*(*i, t*);
- Abundance, namely the number of individuals in the school *N*(*i, t*);
- Spatial location, i.e. the grid cell *c*(*i, t*) where the school is located; and
- Taxonomic identity, i.e. the species *s*(*i*) to which the school belongs.

**Table 1:** Species-specific parameters associated to the evolutionary submodel and the selective mortalities in Ev-OSMOSE

| Symbol | Description | Units | Equations | Source |
| --- | --- | --- | --- | --- |
| Genome structure |  |  |  |  |
| $l_z$ | Number of functional loci for trait $z$ ($z \in \{I_{\max}, m_0, m_1, r\}$) | – | 1 | Assumed |
| $n_{z,l}$ | Number of possible allelic values at functional locus $l$ ($l \in \{1, 2, \dots, l_z\}$) for trait $z$ ($z \in \{I_{\max}, m_0, m_1, r\}$) in the initial population | – | | Assumed |
| $l_b$ | Number of neutral loci | – | | Assumed |
| $n_{b,l}$ | Number of possible allelic identities at neutral locus $l$ ($l \in \{1, 2, \dots, l_b\}$) in the initial population | – | | Assumed |
| $\bar{G}_z(0)$ | Initial mean genotypic value of trait $z$ ($z \in \{I_{\max}, m_0, m_1, r\}$) in the population | Trait unit | 1 | Estimated<sup>1</sup><br>($m_0, m_1, r$) or calibrated<br>($I_{\max}$) |
| $\sigma_{A,z}^2(0)$ | Initial additive genetic variance of trait $z$ ($z \in \{I_{\max}, m_0, m_1, r\}$) in the population | Trait unit | | Calibrated or assumed |
| Trait expression |  |  |  |  |
| $\bar{e}_z$ | Mean expression noise for trait $z$ ($z \in \{I_{\max}, m_0, m_1, r\}$) | Trait unit | | Randomly drawn |
| $\sigma_{e,z}^2$ | Expression noise variance for trait $z$ ($z \in \{I_{\max}, m_0, m_1, r\}$) | Trait unit | | Calibrated or assumed |
| Mortality |  |  |  |  |
| $F_{\max}$ | Maximum instantaneous fishing mortality rate | timestep<sup>-1</sup> | | Calibrated |
| $k_1$ | Instantaneous foraging mortality rate for an individual with an $I_{\max}$ value equal to the initial mean genotypic value of the trait in the population | timestep<sup>-1</sup> | 4 | Calibrated |
| $k_2$ | Exponential slope of the instantaneous foraging mortality | cm<sup>-1</sup> | 4 | Calibrated |

<sup>1</sup> from SMALK data (sex–maturity–age–length key)

**Table 2:** Variables and functions used in the Ev-OSMOSE model.

| Symbol | Description | Units | Equations |
| --- | --- | --- | --- |
| **Entities: Fish schools** |  |  |  |
| Genetic determinism and expression |  |  |  |
| *State variables* |  |  |  |
| $A_{z,l,k}(i)$ | Additive effect of allele $k$ ($k \in \{1,2\}$ as individuals are diploid) at locus $l$ ($l \in \{1,2, \dots, l_z\}$) for trait $z$ ($z \in \{I_{\max}, m_0, m_1, r\}$) of school $i$ | Trait unit | 1 |
| $e_z(i)$ | Phenotypic expression noise for trait $z$ ($z \in \{I_{\max}, m_0, m_1, r\}$) of school $i$ | Trait unit | 2 |
| $b_{l,k}(i)$ | Identity of neutral allele $k$ ($k \in \{1,2\}$ as individuals are diploid) at locus $l$ ($l \in \{1,2, \dots, l_b\}$) of school $i$ | – | |
| *Traits: Emerging individual variables* |  |  |  |
| $G_z(i)$ | Genotypic value of trait $z$ ($z \in \{I_{\max}, m_0, m_1, r\}$) for school $i$ | Trait unit | 1,2 |
| $z(i)$ | Phenotypic value of trait $z$ ($z \in \{I_{\max}, m_0, m_1, r\}$) for school $i$ | Trait unit | 2 |
| $I_{\max}(i)$ | Maximum mass-specific ingestion rate of school $i$ | $g \cdot g^{-\beta} \cdot \text{timestep}^{-1*}$ | |
| $r(i)$ | Gonado-somatic index of school $i$ | – | |
| $m_0(i)$ | Intercept of the maturation reaction norm of school $i$ | cm | |
| $m_1(i)$ | Slope of the maturation reaction norm of school $i$ | $\text{cm} \cdot \text{y}^{-1}$ | |
| *Genetic inheritance: Emerging individual variables* |  |  |  |
| $p_i(t)$ | Probability of school $i$ to be one of the 2 parents of a given new school produced during the breeding season starting at time step $t$ | – | 3 |
| *Ontogenic state* |  |  |  |
| *State variables* |  |  |  |
| $a(i, t)$ | Age of school $i$'s individuals at time step $t$ | y | |
| $w(i, t)$ | Somatic mass of school $i$'s individuals at time step $t$ | g | |
| $g(i, t)$ | Gonadic mass of school $i$'s individuals at time step $t$ | g | |
| *Emerging individual variables* |  |  |  |
| $L(i, t)$ | Total length of school $i$'s individuals at time step $t$ | cm | |
| $m(i, t)$ | Maturity state of school $i$'s individuals at time step $t$ | – | |
| $a_m(i)$ | Maturation age of school $i$'s individuals | y | |
| $w_m(i)$ | Maturation somatic mass of school $i$'s individuals | g | |
| $L_m(i)$ | Maturation length of school $i$'s individuals | cm | |
| $N_{\text{eggs}}(i, t)$ | Total fecundity of school $i$ at first time step $t$ of the breeding season | # | 3 |
| Abundance: *State variable* |  |  |  |
| $N(i, t)$ | Number of individuals in school $i$ at time step $t$ | # | 5 |
| Biomass: *Emerging variables* |  |  |  |
| $B(i, t)$ | Biomass of school $i$ at time step $t$ | g | |
| Spatial Location: *State variable* |  |  |  |
| $c(i, t)$ | Grid cell of school $i$ at time step $t$ | – | |
| Taxonomic identity: *State variable* |  |  |  |
| $s(i)$ | Species to which school $i$ belongs | – | 3 |
| Mortality: *Emerging variables* |  |  |  |
| $M_f(i)$ | Instantaneous foraging mortality rate of school $i$ | timestep<sup>-1</sup> | 4,5 |
| $F(i)$ | Instantaneous fishing mortality rate of school $i$ | timestep<sup>-1</sup> | |
| $\mu_l(i)$ | Instantaneous larval mortality rate of school $i$ | timestep<sup>-1</sup> | |
| $\mu(i)$ | Instantaneous diverse mortality rate of school $i$ | timestep<sup>-1</sup> | |
| **Entities: Fish populations** |  |  |  |
| Abundance: *Emerging population variables* |  |  |  |
| $N(t)$ | Population census size at time step $t$ | # | |
| $B(t)$ | Population biomass at time step $t$ | ton | |
| $C(t)$ | Fishing catches at time step $t$ | ton | |
| Trait distribution: *Emerging population variables* |  |  |  |
| $\bar{G}_z(t)$ | Population genotypic mean of trait $z$ ($z \in \{I_{\text{max}}, m_0, m_1, r\}$) at time step $t$ | Trait unit | |
| $\bar{z}(t)$ | Population phenotypic mean of trait $z$ ($z \in \{I_{\text{max}}, m_0, m_1, r\}$) at time step $t$ | Trait unit | |
| $\sigma_{A,z}^2(t)$ | Population additive genetic variance of trait $z$ ($z \in \{I_{\text{max}}, m_0, m_1, r\}$) at time step $t$ | Trait unit | |
| $\sigma_z^2(t)$ | Population phenotypic variance of trait $z$ ($z \in \{I_{\text{max}}, m_0, m_1, r\}$) at time step $t$ | Trait unit | |
| **Spatial scales and units: grid cells** |  |  |  |
| Spatial coordinates: *State variables* |  |  |  |
| $x(c)$ | Longitude of grid cell $c$ | | |
| $y(c)$ | Latitude of grid cell $c$ | | |
| Physico-chemical factors: *State variables* |  |  |  |
| $pc_k(c, t)$ | Value of physico-chemical factor $k$ of grid cell $c$ at time step $t$ | | |
| $T(c, t)$ | Temperature of grid cell $c$ at time step $t$ | K | |
| $[O_2](c, t)$ | Dissolved O<sub>2</sub> saturation of grid cell $c$ at time step $t$ | % | |
| Biomass of lower trophic levels: *State variables* |  |  |  |
| $B_{LTL}(c, t, j)$ | Biomass of trophic level $LTL\ j$ of grid cell $c$ at time step $t$ | | |

\* $\beta$ is the scaling exponent of maximum ingestion rate and maintenance rate with body mass.

A number of variables further characterizing the individuals of each school emerge from the three first categories of state variables (and thus are not strictly speaking state variables themselves). In terms of trait genetic determinism and expression, the effects of functional loci translate into a genotypic value *G_z_*(*i*) for each evolving trait *z*. Trait phenotypic values result from the influence of both the genotypic value *G_z_*(*i*) and the phenotypic expression noise *e_z_*(*i*). There are four evolving traits in the model, and hence phenotypic values, namely maximum mass-specific ingestion rate *I*_max_(*i*) that determines individuals’ maximum energy uptake from predation, gonado-somatic index *r*(*i*) that determines their energy allocation to somatic growth and reproduction, and two traits that specify their maturation schedule, that is the intercept *m*_0_(*i*) and the slope *m*_1_(*i*) of a deterministic linear maturation reaction norm (Stearns and Koella, 1986). Their evolution allows us to model the evolution of the three life history traits most described in response to fishing-induced evolution (Heino et al., 2015). Schools are also further described by emerging variables such as individuals’ total body length *L*(*i, t*) and their sexual maturity status *m*(*i, t*) that allows distinguishing between juveniles and adults.

Fish schools are distributed on a horizontal spatial grid that is composed of regular cells and that covers the geographical range of the ecosystem represented. A cell *c* is characterized by its spatial coordinates, longitude *x*(*c*) and latitude *y*(*c*), and (i) physical and (ii) biogeochemical variables respectively: (i) the vertically-integrated value of physico-chemical factors *pc_k_*(*c,t*) (such as temperature *T*(*c, t*) or the level of oxygen saturation (%) [*O*_2_](*c*, *t*)) and (ii) the biomass of each lower trophic level group (indexed by *j*) *B_LTL_*(*c, t,j*) that are not explicitly modeled but provided as input to Ev-OSMOSE from coupled hydrodynamic and biogeochemical models.

All schools belonging to the same species form a population and populations of different species form the fish community. Several aggregated population-based metrics can be tracked at the population level such as abundance *N*(*t*), biomass *B*(*t*), fishing catches *C*(*t*) but also the genotypic and phenotypic means 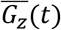 and 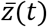 and variances 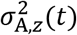 and 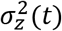 of trait *z* (with *z* ∈ {*I*_max_, *m*_0_, *m*_1_, *r*}.

#### 2.1.2. Design concepts

Ev-OSMOSE relies on a number of well-established concepts and theories and combines them in an original way to describe marine fish biodiversity and its dynamics from the intra-specific - genetic and phenotypic variability - to the inter-specific - taxonomic and trait-based - level. Previous multi-species models of fish communities have been designed to project interspecific biodiversity trajectories under various scenarios considering only ecological dynamics. However, fish populations may also adapt to natural and anthropogenic pressures via phenotypic plasticity and/or evolutionary changes, leading to modifications in their physiology and life-history that could either mitigate or worsen the consequences of these pressures. Ev-OSMOSE has been precisely developed to account for plastic and evolutionary dynamics in fish biodiversity projections by introducing the following elements to the existing OSMOSE model.

Ev-OSMOSE explicitly describes mendelian inheritance of quantitative traits determined by polygenic genotypes according to quantitative genetic principles. The genotypes are composed of a finite number of loci and alleles per locus with effects of heterogeneous amplitude (Soularue and Kremer, 2012), which allows accounting for realistic adaptive and neutral (genetic drift) evolutionary changes and genetic erosion induced by natural and anthropogenic selective pressures. Genetically determined quantitative traits affect individuals’ bioenergetics and sexual maturation processes, which are described with a bioenergetic sub-model.

Individuals’ bioenergetics are described according to a biphasic growth model (Andersen, 2019; Boukal et al., 2014; Quince et al., 2008) in which body mass-dependent energy fluxes are allocated between competing processes—namely maintenance, somatic growth and gonadic growth—thus accounting for physiological trade-offs that constrain both phenotypically plastic and evolutionary responses of life-history traits to selective pressures (Roff, 1992; Stearns, 1992). Moreover, energy fluxes depend on temperature and dissolved oxygen so that metabolic rates follow the oxygen- and capacity-limited thermal tolerance theory (OCLTT; Pörtner, 2001). The details on the bioenergetic sub-model are published in the description of the Bioen-OSMOSE model (Morell et al., 2023).

#### 2.1.3. Emerging properties: fitness, evolution and adaptation

Emergence of most phenomena or characteristics at higher organization levels than the individual one (e.g. population and community spatio-temporal dynamics, population and community age and length structures, species diet) are the same as in the original OSMOSE model.

Phenotypic values of schools’ evolving traits—maximum ingestion rate *I*_max_(*i*), gonado-somatic index *r*(*i*), intercept *m*_0_(*i*) and slope *m*_1_(*i*) of the maturation reaction norm—are entirely determined by their genotype and a randomly drawn expression noise. In contrast, other individual variables or traits at higher integrative levels of organization (hereafter named “emerging variables”: somatic mass *w*(*i,t*), length *L*(*i,t*), gonadic mass *g*(*i,t*) and thus fecundity *N_eggs_*(*i,t*), maturation age *a*_m_(*i*) and somatic mass *w*_m_(*i*) or length *L*_m_(*i*) at maturation) emerge from the combination of evolving traits’ values, energy intake from length-based opportunistic predation and physiological or plastic responses of bioenergetics to ambient sea water temperature and dissolved oxygen concentration (Fig. 2).

The distribution of genotypic and phenotypic values of evolving traits at the population level are fully prescribed initially by the values of the parameters describing genetic variability, namely the initial genotypic mean value 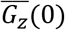 and the initial additive genetic variance 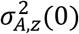, and the expression noise distribution, namely the expression variance 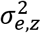, for a given trait *z*. As the simulation progresses, these distributions are affected by the processes of natural, fishing-induced and climate-induced selection and genetic drift so that their changes through time describe emerging evolutionary trajectories. Temporal changes in the phenotypic distribution of emerging variables result from both the trajectories of the underlying evolving traits and phenotypically plastic responses to available food and ambient physico-chemical conditions.

The evolving trait values are variable in the populations and confer advantages or disadvantages in terms of survival and reproductive success relative to different pressures, notably predation, fishing, and climate changes. Therefore, Darwinian fitness, that governs the above-mentioned evolutionary trajectories together with genetic variability, emerges naturally from the modelled processes of mortality and reproduction. In consequence, populations may adapt to predation, fishing and climate change through evolution.

#### 2.1.4. Initialization

For each species, the initial pool of allele values present in the population for each functional or neutral locus is randomly drawn from a prescribed distribution (see section 2.1.6.1 Genetic structure for details). The system starts with no school in the domain and is initialized by releasing eggs for every species during specific reproductive season time steps. For a given species, this seeding process stops when there is at least one mature individual in the population. The eggs are grouped in super-individuals, representing schools that are distributed spatially according to their habitat maps. During the spin-up period (until the system reaches an equilibrium), for each new school of eggs, a diploid genotype is randomly drawn from the functional and neutral pools of alleles at each locus. The mendelian transmission of genotype from parents to offspring starts at the end of the spin-up period.

#### 2.1.5. Input

Ev-OSMOSE does not model oceanographic physical and chemical processes, but it is forced by spatially and temporally varying fields of temperature (°C) and oxygen (% of saturation) from coupled regional physical and biogeochemical models, data time series or from the regional downscaling of earth system model outputs. As for the OSMOSE model, biomass prey fields are also used as input to provide LTL.

#### 2.1.6. Genetic sub-model

The genetic sub-model introduces a source of intra-specific variability of the quantitative traits describing the individual life history, through additive genetic variance 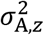 and expression variance 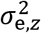, and parental gene inheritance. The genotypic values of the four heritable traits—maximum mass-specific ingestion rate *I*_max_, gonado-somatic index *r*, intercept *m*_0_ and slope *m*_1_ of linear maturation reaction norm—result from the expression of the corresponding functional loci. Neutral loci have no effect on individuals’ phenotype: their evolution is the result of random drift. Following temporal changes in genetic variability at neutral loci is thus a way to assess genetic drift. Hereafter, the genetic sub-model is described for any of the four evolving traits, generically denoted *z*.

##### 3.1.6.1. Genetic structure

The genetic structure is described by a polygenic multi-allelic model with finite numbers of loci and alleles for both the functional and neutral parts of the genome. The value of trait *z* thus results from the expression of *l_z_* functional loci, each of which has a pool of *n_z,l_* (with *l* ∈ { 1,2 ⋯, *l_z_*}) possible alleles in the initial population characterized by *n_z,l_* allelic values. Following classical quantitative genetics (Lynch and Walsh, 1998), we assume that the genotypic values *G_z_*(*i*) of trait *z* in the population initially follow a normal distribution 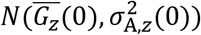 with 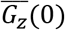 the initial genotypic mean and 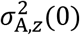 the initial additive genetic variance. It follows (see justification in the next section) that the *n_z,l_* allelic values of locus *l* initially present in the population are randomly drawn from a normal distribution 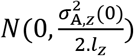 (Soularue and Kremer, 2012). This allelic model defines allelic values as deviations around the initial genotypic mean 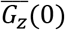 of the population and allows for heterogeneous allelic values across loci coding for the same trait, many of them with minor effects and a few ones with major effects.

Similarly, the neutral part of the genome is described by *l_b_* neutral loci, each of which has a pool of *n_b,l_* (with *l* ∈ { 1,2, ⋯, *l_b_*}) possible alleles in the initial population characterized by their allelic values with no effect on evolving traits. The *n_b,l_* allelic identities of locus *l* initially present in the population are randomly drawn from a discrete uniform distribution with probability mass function 1/*n_b,l_*.

##### 3.1.6.2. Traits’ genetic determinism and expression

The two additive effect allele values *A*_*z,l*,1_(*i*) and *A*_*z,l*,2_(*i*) at a functional locus *l*(*l* ∈ {1,2, ⋯, *l_z_*}) coding for trait *z*(*i*) of diploid individual *i* can each take one allelic value among the *n_z,l_* possible versions in the population. Alleles act additively at and between loci. Since allelic values describe deviations around the mean genotypic value of trait *z*, the genotype value *G_z_*(*i*) for trait *z*(*i*) in school *i* is thus the sum of the initial genotypic mean 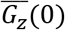 of the trait for the population and of the two allelic values *A_z,l,k_*(*i*) at each locus *l* coding for the trait of interest.

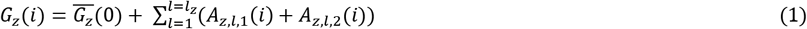

Given the normal distribution additive property and that the initial distributions 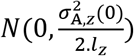 of allelic values in the population are independent between loci, the initial distribution of genotypic values *G_z_*(*i*) in the population thus follows a normal distribution 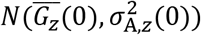. At later time steps *t*, the processes of selection, drift and inheritance will modify this distribution in terms of its mean 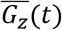 and its variance 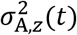 but also potentially in terms of its shape as it is not constrained to remain normally distributed.

In Ev-OSMOSE, part of the phenotypic expression of emerging variables (e.g., somatic mass *w*(*i,t*), gonadic mass *g*(*i,t*), length *L*_m_(*i*) at maturation) is due to the bioenergetic responses to conditions faced by an individual: the available food, the temperature and the oxygen concentration in the environment during the entire individual life cycle. In contrast, the four evolving traits (maximum mass-specific ingestion rate *I*_max_, gonado-somatic index *r*, intercept *m*_0_ and slope *m*_1_ of linear maturation reaction norm) describe underlying individual characteristics whose phenotypic expression does not depend on these “macro-environmental” conditions. Yet, the phenotypic expression of evolving traits will also be affected by dominance and recessivity of alleles at the same locus and epistasis between loci, which are not modeled explicitly in the present genetic model, as well as by “micro-environmental” variations capturing the potentially unaccounted effects of individuals’ internal environment or external micro-environment (Lynch and Walsh, 1998). These sources of phenotypic variability for evolving trait *z* are implicitly represented by an expression noise *e_z_*(*i*) randomly drawn from a normal distribution 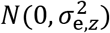 at the individual’s birth and added to the genotypic value of its trait *z*. The phenotypic value of evolving trait *z*(*i*) for the school *i* is then

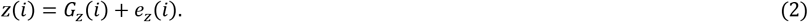

##### 3.1.6.3. Genetic inheritance

Both functional and neutral loci follow Mendelian inheritance under sexual reproduction. Reproduction is panmictic, which means that all sexually mature individuals can contribute to mating pairs of parents irrespective of their location and phenotype. If a new school is created at time step *t*, its two parents are randomly drawn from a multinomial distribution *M*(2,**p**(*t*)) for 2 trials with a probability vector **p**(*t*) composed of as many elements *p_i_*(*t*) as there are schools in the population. The *i*th element *p_i_*(*t*) is defined as the relative fecundity of school *i* in the population at time step *t*,

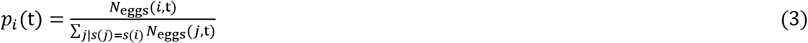

with *N*_eggs_(*i,t*) the fecundity of school *i* and 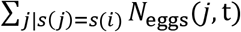 the total fecundity of the species *s*(*i*) population at time step *t*.

For each selected parental school, haploid gametes are assembled by randomly drawing one of the two alleles at each locus to represent allelic segregation during meiosis. This is done under the assumption of independence between loci, so that alleles recombine freely. New schools receive at each functional and neutral locus one allele from both chosen parents by randomly picking a haploid gamete for each of them.

#### 2.1.7. Bioenergetics and life-history sub-model

The four evolving traits of a school *i*—*I*_max_, *r*, *m*_0_ and *m*_1_—together with its age *a*(*i, t*) and somatic mass *w*(*i,t*) determine its bioenergetics and life-history processes, namely somatic and gonadic growth, maturation, reproduction and mortality. The detailed description of the bioenergetics fluxes is provided in Morell et al. (2023). A general description of the bioenergetic fluxes is presented hereafter as well as their linkages with the four evolving parameters (Fig. 3).

**Figure 3:**
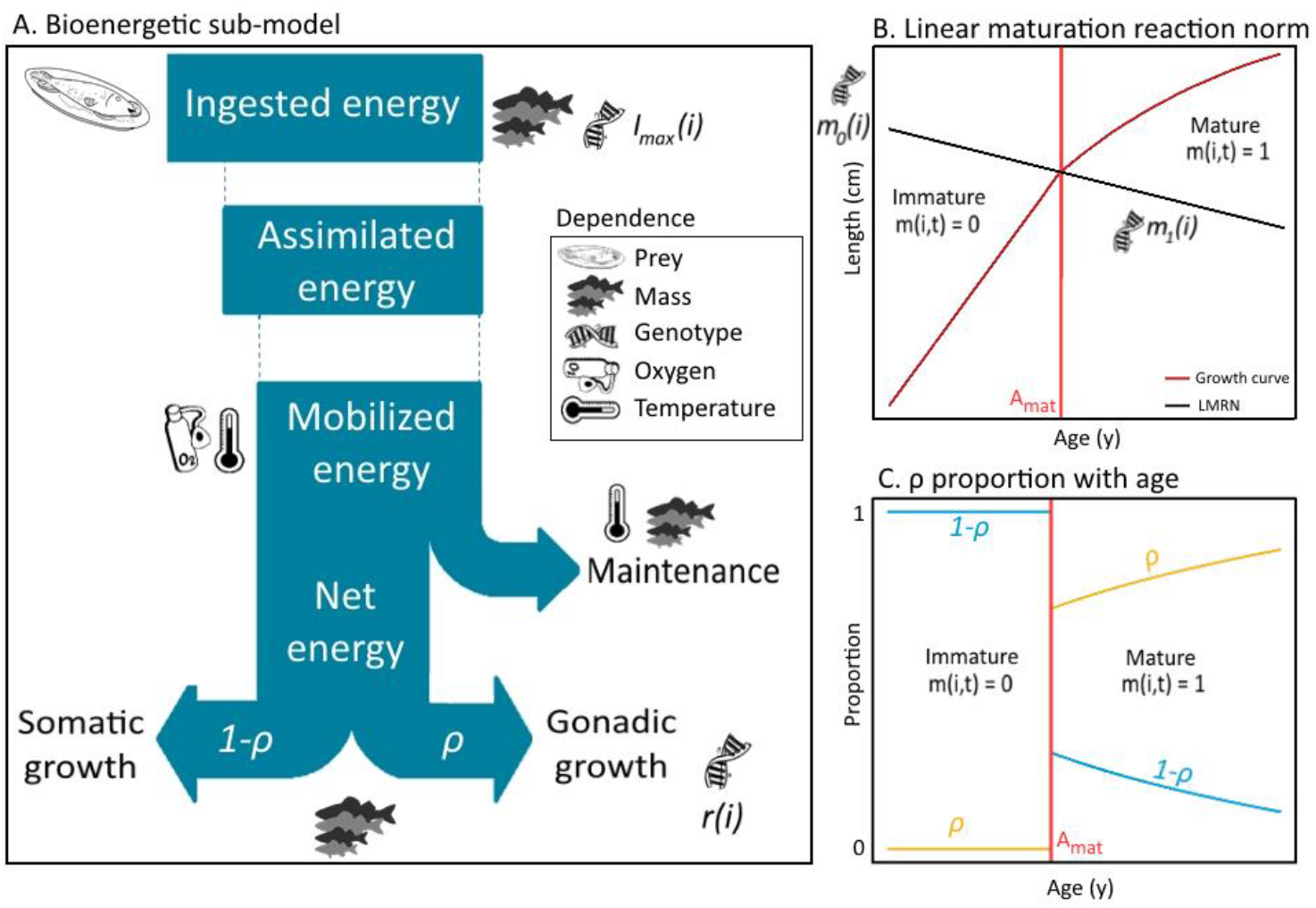
Bioenergetic sub-model fluxes from the ingestion to the tissue growth, namely somatic and gonadic growth (A). The flux dependences to biotic (individual genotype, available prey and somatic mass) and abiotic (temperature and oxygen) variables are specified with pictograms. Four parameters are prone to evolve: the maximum mass-specific ingestion rate *I*_max_ whose evolution impacts the ingested energy and downstream fluxes, the intercept *m*_0_ and the slope *m*_1_ of the linear maturation reaction norm (LMRN) (B) whose evolutions impact the maturation process, and the gonado-somatic index *r* whose evolution impacts the slope of the proportion of net energy *ρ* allocated to gonadic growth after maturity (C) and thus impacts the growth-reproduction tradeoff. The LMRN (B) models all the age-length combinations at which an individual can become mature.

##### 3.1.7.1. General principles

Individual life history emerges from underlying bioenergetic fluxes which are described according to a biphasic growth model (Fig. 3) (Andersen, 2019; Boukal et al., 2014; Quince et al., 2008). The body mass-dependent energy fluxes are allocated according to physiological tradeoffs between competing processes: maintenance, somatic growth and gonadic growth. The sexual maturation of individuals relies on the concept of maturation reaction norms that depicts how the process of maturation responds plastically to variation in body growth (Heino et al., 2002; Stearns and Koella, 1986). This combination of processes mechanistically describes how somatic growth, sexual maturation and reproduction emerge from energy fluxes sustained by food intake resulting from opportunistic length-based predator-prey interactions.

On top of the biphasic growth model, individuals’ energy mobilization and maintenance energetic costs depend on dissolved oxygen saturation and temperature so that the resulting metabolic rate (the net energy available for new tissue production) and thus somatic and gonadic growth vary with these abiotic parameters in a way that conforms to the oxygen- and capacity-limited thermal tolerance theory (OCLTT; Pörtner, 2001) and more generally to thermal performance curves (TPC; Angilletta, 2009). The equations underlying the bioenergetic sub-model and especially the plastic responses to dissolved oxygen saturation and temperature are not developed hereafter as they are fully described in a previous paper (Morell et al., 2023). As Ev-OSMOSE models the evolution of bio-energetic process traits underlying the life history, we propose a simplified description of the bioenergetic processes that are essential to understand the role of the traits, also illustrated with Fig. 3.

##### 3.1.7.2. Fluxes description: from the ingestion of energy to tissue growth

The bioenergetic fluxes are summed up in Fig. 3A. The most upstream flux is the ingestion of energy. The ingested energy follows a Type 1 functional response to prey biomass: it increases linearly with the amount of prey biomass that is spatiotemporally co-occurring with the feeding school, until it reaches a maximum that increases with individual somatic mass, corresponding to the satiety state level. The predator-prey co-occurrence depends on the spatial distributions of the prey (other HTL schools and forcing LTL prey fields) and of the feeding schools.

A constant portion of the ingested energy is assimilated. The portion which is not assimilated is lost due to excretion and feces egestion. A portion of assimilated energy is then mobilized. The mobilized energy pays internal processes, i.e, growth of the somatic and gonadic tissues and maintenance in our framework.

The portion of assimilated energy that is mobilized depends on temperature and oxygen. The mobilized energy rate fuels all metabolic processes starting in priority with the costs of maintenance of existing tissues. The maintenance rate increases with temperature and with somatic mass. The difference between mobilized energy and maintenance is called net energy for new tissue production. The net energy is then fully allocated to the growth of the somatic compartment before maturation and it is shared between growth of the somatic and gonadic compartments after maturation. The increase of the somatic compartment implies growth in length and mass. The energy allocated to the gonadic compartment is used during the breeding season to produce eggs.

The maturation process is modeled by a deterministic linear maturation reaction norm (LMRN) that represents all the age-length combinations at which an individual can become mature (Stearns and Koella, 1986; Stearns, 1992) (Fig. 3B). In this framework, individuals become sexually mature when their growth trajectory in terms of body length intersects the LMRN. The mature state *m*(*i, t*) is 0 for immature individuals and 1 for mature individuals.

#### 2.1.8. Mortality

The mortality sub-model is described in the Supporting Information in Morell et al. (2023). To sum up the mortality process, a school *i* faces different sources of mortalities at each time step, namely predation mortality caused by other schools (emerging), starvation mortality (emerging), fishing mortality *F*(*i*), larval mortality *μ_l_*(*i*) and diverse other natural mortalities *μ*(*i*) (i.e. senescence, diseases, and non-explicitly modeled predators). An additional foraging mortality is modeled in Ev-OSMOSE. This mortality describes the additional mortality due to foraging for prey. Each time step *t* is subdivided into multiple sub-time steps *dt* within which the different mortality sources impact a school *i* in a random order so as to simulate the simultaneous nature of these processes (see http://documentation.osmose-model.org/ for more details). Hereafter, we detail the mortality that represents the main selective pressures and/or important evolutionary tradeoffs in our framework.

Organisms face a trade-off between foraging activity and mortality (Mangel, 2003) because more active foraging implies a higher exposure to predation, unfavorable conditions (e.g., triggering diseases) and/or increased oxidative stress. Assuming that variation in mass-specific maximum ingestion *I*_max_ rate results from variation in foraging activity, this trade-off is modeled by including a foraging mortality that increases with the mass-specific maximum ingestion rate *I*_max_ and thus when foraging activity is more intense. The instantaneous foraging mortality rate experienced by school is defined as follows:

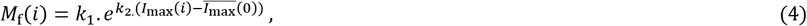

with *k*_1_ the foraging mortality that would face an individual *i* if it had an *I*_max_(*i*) value equal to the initial mean genotypic value of the trait 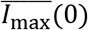 in the population and *k*_2_ the exponential slope translating the change of foraging activity linked to a deviation of *I*_max_(*i*) from 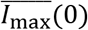 into an multiplicative factor of the trade-off’s strength. Change in the number of individuals in school *i* due to foraging mortality during sub-time step *dt* is then obtained as:

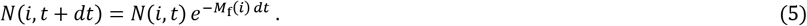

Fishing mortality is a major evolutionary pressure on marine populations due to total mortality increase and length selectivity. In the model, fishing mortality can be discretized per length class, i.e., a parameter of fishing mortality per species per length class can be used to realistically model the fishing process. The highest fishing mortality rate across length classes of a species is called *F_max_*.

Predation-induced mortality is an explicit stochastic length-dependent process that emerges from the spatial co-occurrence between predators and prey, and the predators’ ingestion process. The predation mortality applied to school *i* is simply the sum of the biomass losses due to the ingestion of all predator schools *j* with suitable body length, and present in the same grid cell *c*(*i,t*) at subtime step *dt*. From length-dependent interactions emerge a realistic selective predation pressure that decreases with fish length.

Starvation mortality occurs when an individual cannot cover its energetic maintenance needs, i.e. when net energy is negative. If the energy reserve, provided by gonads, is not sufficient to cover the maintenance needs, the school undergoes an energetic deficit and faces starvation mortality proportionally to its energy deficit. In our model, starvation mortality increases in response to climate change due to rising temperature, deoxygenation or decrease in food availability. The increase of total mortality through increased starvation mortality is expected to accelerate life cycle similarly to what is expected under fishing pressure (Waples and Audzijonyte, 2016).

### 2.2. The North Sea ecosystem application: Ev-OSMOSE-NS

#### 2.2.1. Application presentation

The Bioen-OSMOSE model, i.e without the evolutionary sub-model, was applied to the North Sea ecosystem and published in Morell et al. (2023) and summed up in Fig. 4. The model domain is delimited by the Norwegian Trench in the north east and includes the eastern English Channel. The grid is regular with cells of 0.25° x 0.5° (632 sea cells). The Ev-OSMOSE-NS, i.e. including the evolutionary sub-model, models 15 fish species (Fig. 4). The configuration represents a mean steady state of the ecosystem for the period 2010-2019. The full description of the parameterization of the 15 fish species is provided in Morell et al., (2023). Hereafter, we detail the parameterization of the new evolutionary sub-model and the calibration that was performed with this new sub-model.

**Figure 4:**
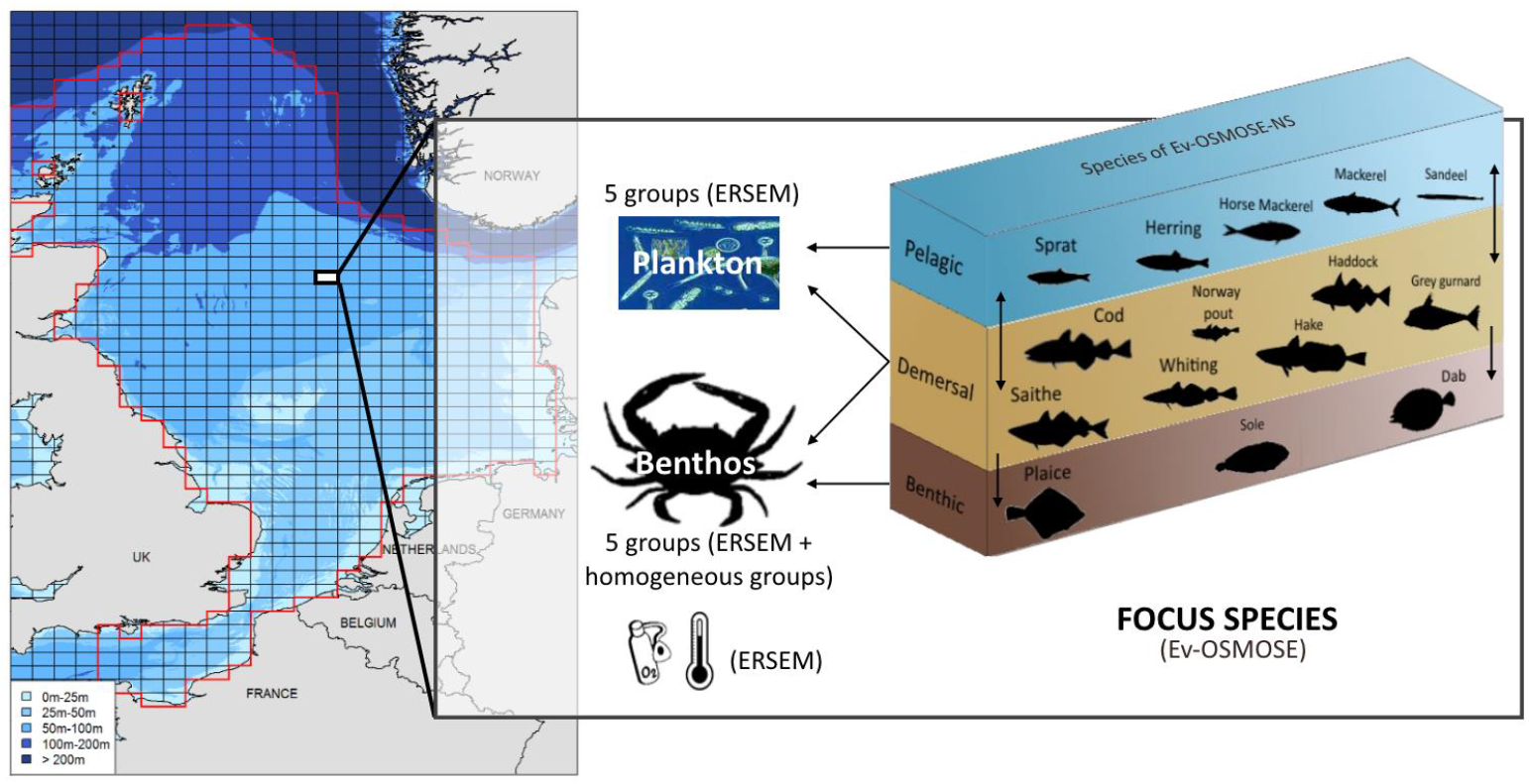
Representation of the Ev-OSMOSE-NS model applied to the North Sea and the Eastern Channel. Fifteen focus species are explicitly modeled. Outputs from the coupled POLCOMS-ERSEM model force Ev-OSMOSE: temperature, oxygen, and the biomass of 8 LTL plankton and benthic groups. Two homogeneous benthic groups are added to model large benthic prey.

#### 2.2.2. Parameterization of the evolutionary sub-model

For each species and each evolving trait, the required parameters in the evolutionary sub-model are: the initial mean genotypic value 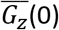, the initial additive genetic variance 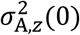, the expression noise variance 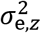, the number of functional loci *l_z_* and the number of allelic values *n_z, l_* for each of them. It necessitates in addition determining values for the foraging mortality coefficients *k*_1_ and *k*_2_. In this first application of the Ev-OSMOSE modelling framework, neutral loci were not activated, but values for the number of neutral loci *l_b_* and the number of allelic identities *n_b,l_* for each of them are also required otherwise.

The mean initial genotypic value 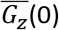 of a trait is by definition equal to the mean phenotypic value of the trait in the population as expression noise and allelic values are centered around 0. The initial mean genotypic/phenotypic values of the traits were thus fixed at the value estimated for the Bioen-OSMOSE-NS configuration (Morell et al. 2023), except for the mean value of *I*_max_ that was calibrated *de novo* for Ev-OMOSE-NS (see next section “Model calibration”).

The initial additive genetic variance 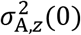 and the expression noise variance 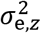 were estimated according to the following procedure. Given additivity and independence of the genetic and micro-environmental effects on the phenotypic value of a trait (equation 2), the phenotypic variance of a trait is the sum of additive genetic variance and expression noise variance 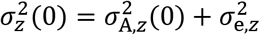. Heritability is defined as the proportion of phenotypic variance due to additive genetic variance, 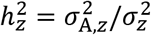, and is typically around 0.2 for life-history traits of vertebrates and ectotherms (Mousseau and Roff, 1987). Given a certain trait phenotypic variance 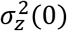 (that can be estimated from field data see below), initial additive genetic variance and expression noise variance can then be estimated as 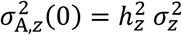 and 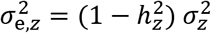.

The phenotypic variances 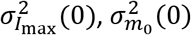, and 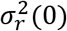 were estimated from variability in length-at-age and maturation for each species using SMALK data (see details in Supporting Information A). For the sake of simplicity, the phenotypic variance of the slope of the LMRN *m*_1_, 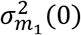, was fixed to 0. This assumption implies that the slope of the LMRN *m*_1_ cannot evolve (if there is no phenotypic variance, there is no additive genetic variance), all the maturation variance is explained by the population phenotypic variance of *m*_0_, 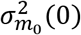, and that the mean maturation length variance is constant at any age (see Supporting Information A2). This is justified by the fact that (i) the first order term in empirically documented evolutionary changes in maturation reaction norm is explained by a change of its intercept *m*_0_ (e.g. Marty et al., (2014) for North Sea gadoids) so that evolution of the slope *m*_1_ can be neglected in first approximation and (ii) population variance in maturation age and length can be correctly approximated by variance in the LMRN intercept only.

In the simulations, the evolution of two out of the three traits with non-zero phenotypic variance was activated, i.e., the genotypic variance was set different from 0, with a heritability of 0.2, for these traits: the gonado-somatic index *r* and the intercept of the LMRN *m*_0_. The evolution of *I*_max_ was not activated because the available data were not suitable to estimate the trade-off between foraging mortality and ingestion and the resulting evolutionary trends would have been subject to caution. However, the choice was made to keep phenotypic variance of *I*_max_ included as it determines directly phenotypic variance in juvenile growth, which is one of the most variable traits in fish. In terms of sources of variance, this assumption means that all the phenotypic variance of *I*_max_ is explained by the expression noise only, 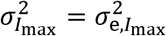. The values of the expression noise variances and the additive genetic variances used for the simulations are given in Table 3.

**Table 3:** Micro-environmental noise and genotypic variances of process traits in Ev-OSMOSE-NS. The sum of these variances is the total phenotypic variance of each trait.

| Species | $I_{\max}$ | | $r$ | | $m_0$ | | $m_1$ | |
| --- | --- | --- | --- | --- | --- | --- | --- | --- |
| | $\sigma_{e,I_{\max}}^2$ | $\sigma_{A,I_{\max}}^2(0)$ | $\sigma_{e,r}^2$ | $\sigma_{A,r}^2(0)$ | $\sigma_{e,m_0}^2$ | $\sigma_{A,m_0}^2(0)$ | $\sigma_{e,m_1}^2$ | $\sigma_{A,m_1}^2(0)$ |
| Herring (*Clupea harengus*) | 0.09 | 0 | 0.02 | 4.85e-03 | 14.07 | 3.52 | 0 | 0 |
| Mackerel (*Scomber scombrus*) | 0.12 | 0 | 0.02 | 1.17e-02 | 9.24 | 2.31 | 0 | 0 |
| Sandeel (*Ammodytes* spp) | 0.12 | 0 | 0.02 | 6.00e-03 | 2.33 | 0.58 | 0 | 0 |
| Sprat (*Sprattus sprattus*) | 0.03 | 0 | 0.01 | 6.38e-03 | 3.68 | 0.92 | 0 | 0 |
| Norway pout (*Trisopterus esmarkii*) | 0.06 | 0 | 0.01 | 6.00e-03 | 30.33 | 7.58 | 0 | 0 |
| Plaice (*Pleuronectes platessa*) | 0.12 | 0 | 0.02 | 3.09e-03 | 45.87 | 11.47 | 0 | 0 |
| Sole (*Solea solea*) | 0.28 | 0 | 0.06 | 1.07e-02 | 21.21 | 5.3 | 0 | 0 |
| Saithe (*Pollachius virens*) | 0.08 | 0 | 0.02 | 3.66e-03 | 180.16 | 45.04 | 0 | 0 |
| Cod (*Gadus morhua*) | 0.63 | 0 | 0.13 | 2.57e-03 | 283.56 | 70.89 | 0 | 0 |
| Haddock (*Melanogrammus aeglefinus*) | 0.36 | 0 | 0.07 | 6.00e-03 | 36.22 | 9.05 | 0 | 0 |
| Horse Mackerel (*Trachurus trachurus*) | 0.48 | 0 | 0.1 | 4.48e-03 | 6.26 | 1.57 | 0 | 0 |
| Whiting (*Merlangius merlangus*) | 0.35 | 0 | 0.07 | 7.31e-03 | 11.54 | 2.89 | 0 | 0 |
| Dab (*Limanda limanda*) | 0.12 | 0 | 0.02 | 6.00e-03 | 12.77 | 3.19 | 0 | 0 |
| Grey gurnard (*Eutrigla gurnardus*) | 0.22 | 0 | 0.04 | 6.00e-03 | 12.24 | 3.06 | 0 | 0 |
| Hake (*Merluccius merluccius*) | 0.4 | 0 | 0.08 | 6.00e-03 | 108.41 | 27.1 | 0 | 0 |

The number of functional loci *l_z_* and alleles per locus *n_z,l_* were fixed to 10 and 7, respectively, based on experience from previous monospecific eco-genetic models (Marty et al. 2015) and analogy with the order of magnitude of the number of allelic values typically observed for neutral markers in fish such as microsatellites (e.g. Poulsen et al., 2006). These values also insured obtaining an initial normal distribution of the traits in the population.

#### 2.2.3. Model calibration

The Bioen-OSMOSE-NS configuration detailed in Morell et al., (2023) was calibrated to obtain estimates for unknown parameters, using maximum likelihood estimation based on an evolutionary optimization algorithm adapted to high-dimensional parameter space that is available in the calibraR R package (Oliveros-Ramos and Shin, 2016). The algorithm explores the space of unknown parameters so as to maximize the likelihood obtained by comparing model outputs to observed data.

The addition of a new evolutionary sub-model to the North Sea configuration modifies the simulation outputs of the model, notably by introducing interindividual variability through phenotypic variance, and thus the Ev-OSMOSE-NS model needed to be calibrated anew to re-estimate the same unknown parameters as in Bioen-OSMOSE-NS. The estimations of the parameters obtained from the calibration of Bioen-OSMOSE-NS were used as initial guesses to speed up the calibration process. The calibration of the Ev-OSMOSE-NS model is an ‘ecological fit’ to ecological data using a model version with phenotypic variability but without genotypic transmission. The data used to calibrate Ev-OSMOSE-NS are fisheries landings (ICES, 2019a), length-at-age from scientific surveys from ICES database (NS-IBTS-Q1, ICES DATRAS 2022) and estimated biomasses for assessed species (ICES, 2016, 2018a, 2018b, 2018c, 2019b). The calibration is performed for an average state of the ecosystem for the period 2010-2019 by using observed data collected over the period as target values (Supporting Information B). For each species, the estimated parameters are the larval mortality rate *μ_l_*(*i*), the mean maximum ingestion rate *I*_max_, the maximum fishing mortality rate *F_max_*, and the additional mortality rate *μ*(*i*). A parameter per LTL group named coefficient of accessibility of fish is also estimated. The new estimation of these parameters for the Ev-OSMOSE-NS model is given in Table 4 for species parameters and Table 5 for LTL parameters. Due to limited data on the relationship between foraging behavior, predation mortality and growth rate, the coefficients *k*_1_ and *k*_2_ for the trade-off between *I*_max_ and foraging mortality were manually tuned and not calibrated through maximum likelihood estimation. They were fixed so that foraging mortality for each species was on average (i.e. when accounting for phenotypic variance in *I*_max_) equal to 0.05 and the slope *k*_2_ was manually tuned (and hence *k*_1_ adjusted to maintain the average at 0.05) to obtain evolutionary trends in *I*_max_ that were within reasonable ecological limits (results not shown obtained by activating evolution for *I*_max_ contrary to the simulations presented here). Values of the two coefficients are also given in Table 4.

**Table 4:**
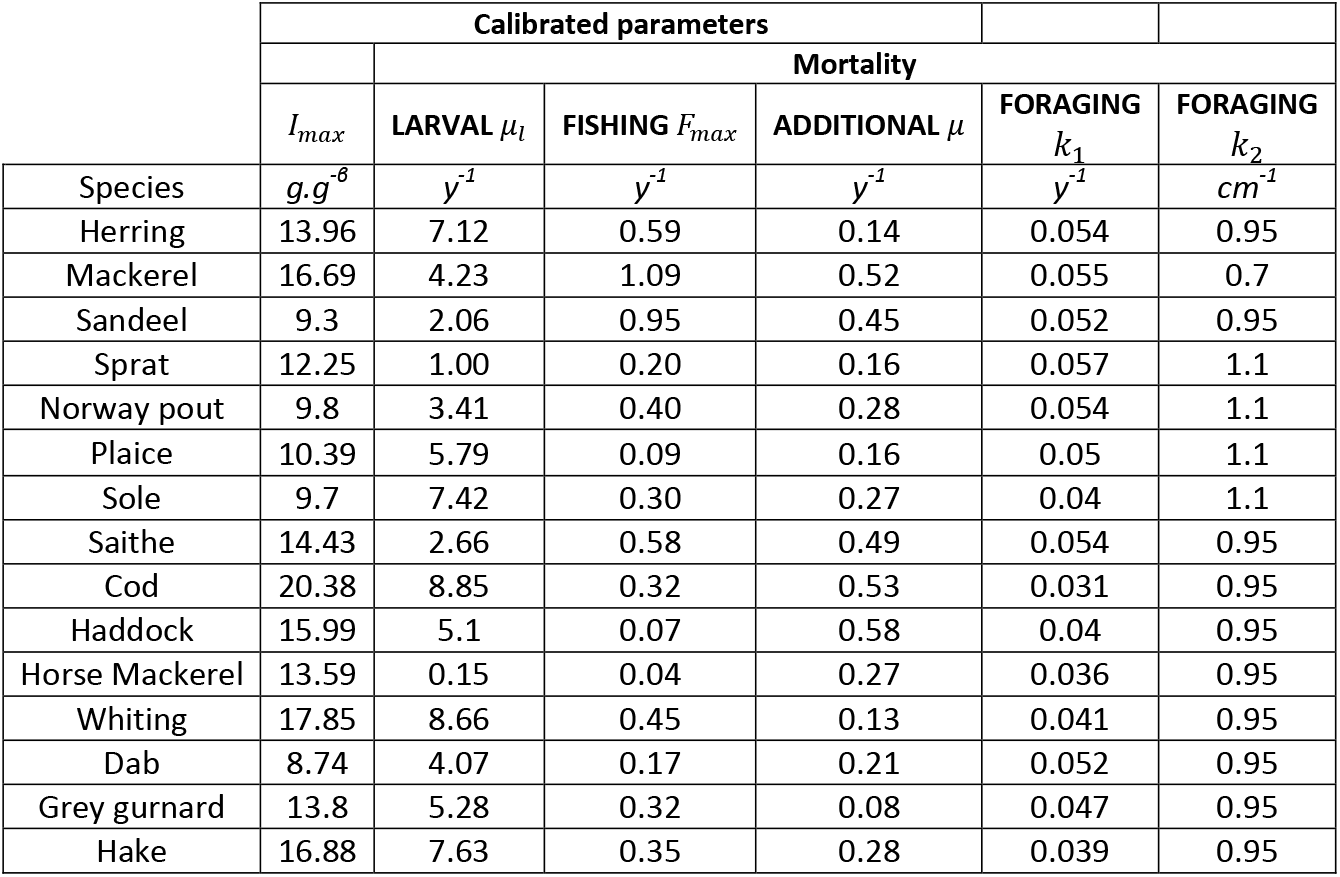
Calibrated species parameters for the 15 fish species in Ev-OSMOSE-NS.

**Table 5:**
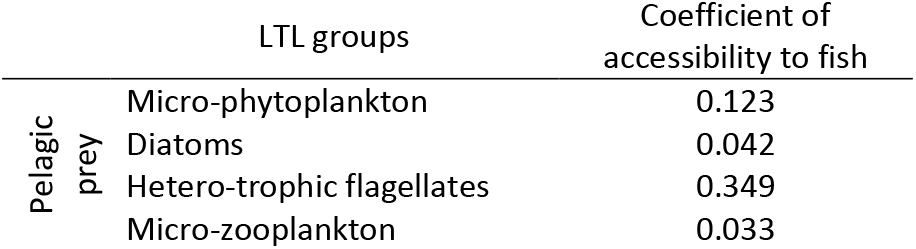
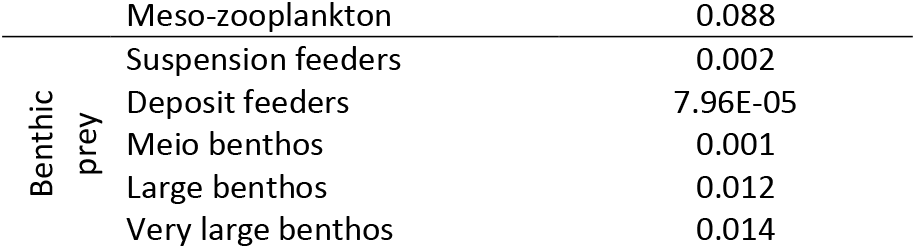
Calibrated coefficients of accessibility of fish to low trophic level (LTL) groups in Ev-OSMOSE-NS.

The calibrated configuration is run for 100 years. The first 50 years is the spin-up period, a period during which the system stabilizes. The years from 50 to 70 constitute the reference stable state of the simulated system without evolution. The maturation and size-at-age outputs from this period are presented in the results. Mendelian transmission is activated on year 70. The transmission results presented hereafter are for the years after the Mendelian transmission activation. 28 replicates of the model are run with the same parameterization to account for Ev-OSMOSE-NS stochasticity.

## 3. Results

The NS configuration has already been calibrated and evaluated in a version without genotypic and phenotypic variance (Bioen-OSMOSE-NS, Morell et al., 2023). To avoid redundancy in this paper, the indicators used to evaluate the ecological validity of the configuration are in Supporting Information B. Since the model’s originality lies in how it includes phenotypic and genotypic variance, indicators demonstrating the model’s capacity to replicate realistic emergent variability received particular attention (see Section 3.1). Considering Bioen-OSMOSE-NS as the reference configuration, we explored for which aspects the new developments in Ev-OSMOSE improve the realism of the model predictions. In consequence, we present how the maturation and length-at-age outputs of Bioen-OSMOSE-NS (Morell et al., 2023) differ from those produced by Ev-OSMOSE-NS and whether Ev-OSMOSE better fits observed data. The ability of the model to account for evolutionary responses that correctly respond to selective pressures is illustrated by the transmission of genotypic values between parental pools and new born cohorts.

### 3.1. Emerging phenotypic variability

#### 3.1.1. Maturation

A comparison of Ev-OSMOSE-NS simulation outputs for maturity ogives with Bioen-OSMOSE-NS (Morell et al., 2023) outputs and observed data can indicate whether taking into account phenotypic variance in process traits improves model realism, especially as maturity ogives were not used as targets for calibration. The maturation process can be assessed with two types of Ev-OSMOSE outputs: (i) the mean maturation age or length and (ii) the variance of the maturation age or length. The slopes of the related maturity ogives can be used to visually assess if the simulated variance better fits the observations.

Compared to Bioen-OSMOSE-NS, Ev-OSMOSE-NS provides a better representation of mean age at maturity for haddock, hake, herring, plaice and sole (closer to observed mean ages at maturity), a similar one for saithe and whiting, but a worse one for cod, grey gurnard, Norway pout and sprat (vertical lines, Fig. 5A). Ev-OSMOSE-NS outputs reproduce better observed variance in mean age at maturity for all species except sprat and mackerel (curves, Fig. 5A). The simulated mackerel ages at maturity fail to reproduce a credible shape for the age-based maturity ogive.

**Figure 5:**
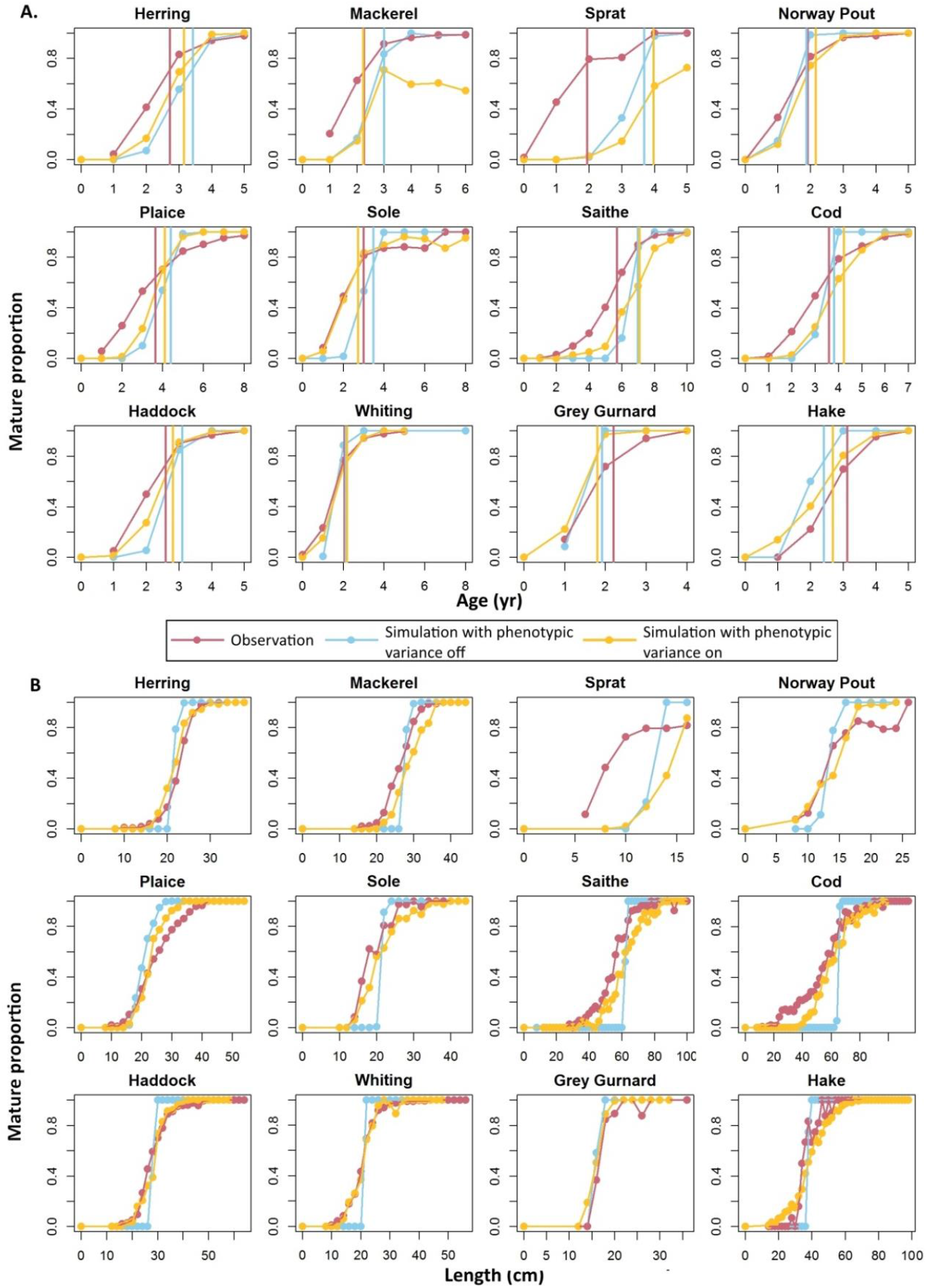
Age- (A) and length-based (B) maturity ogives per species for observed (red), simulated without (blue) and simulated with phenotypic variance (yellow) individual data for species for which empirical maturation data are available. Results are shown for the species for which there is enough data to estimate and plot the observed age and length maturity ogives. Age data are yearly grouped and length data are grouped by 2-centimeter classes. The vertical lines are the mean ages at maturation (A) computed as 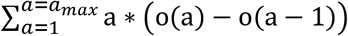 with o(a) the proportion of mature individuals at age a. The mean length at maturation is not represented. Some observed length maturity ogives are not strictly increasing and do not allow a reliable estimation of the mean maturation length.

The evaluation of the model’s maturation outputs is complemented with the length-based maturity ogives (Fig. 5B). Those simulated with Ev-OSMOSE-NS show a much better visually fit to observed ones in terms of both mean and variance of lengths at maturity for all species except sprat. The fit to data is particularly good for haddock, herring and whiting length ogives.

#### 3.1.2. Length-at-age

The evaluation of the model on the simulated lengths-at-ages is performed in a similar way to the maturation indicators: we first inspect the shape of the length-at-age curves (Fig. 6) and we also calculate the sum of squared errors (SSE) between the simulated and observed means and standard deviations of length at different ages (Fig. 7). We chose the SSE of the standard deviations as an indicator of the goodness of fit for length variability at age, because the SSE of the variances would overly highlight outliers.

**Figure 6:**
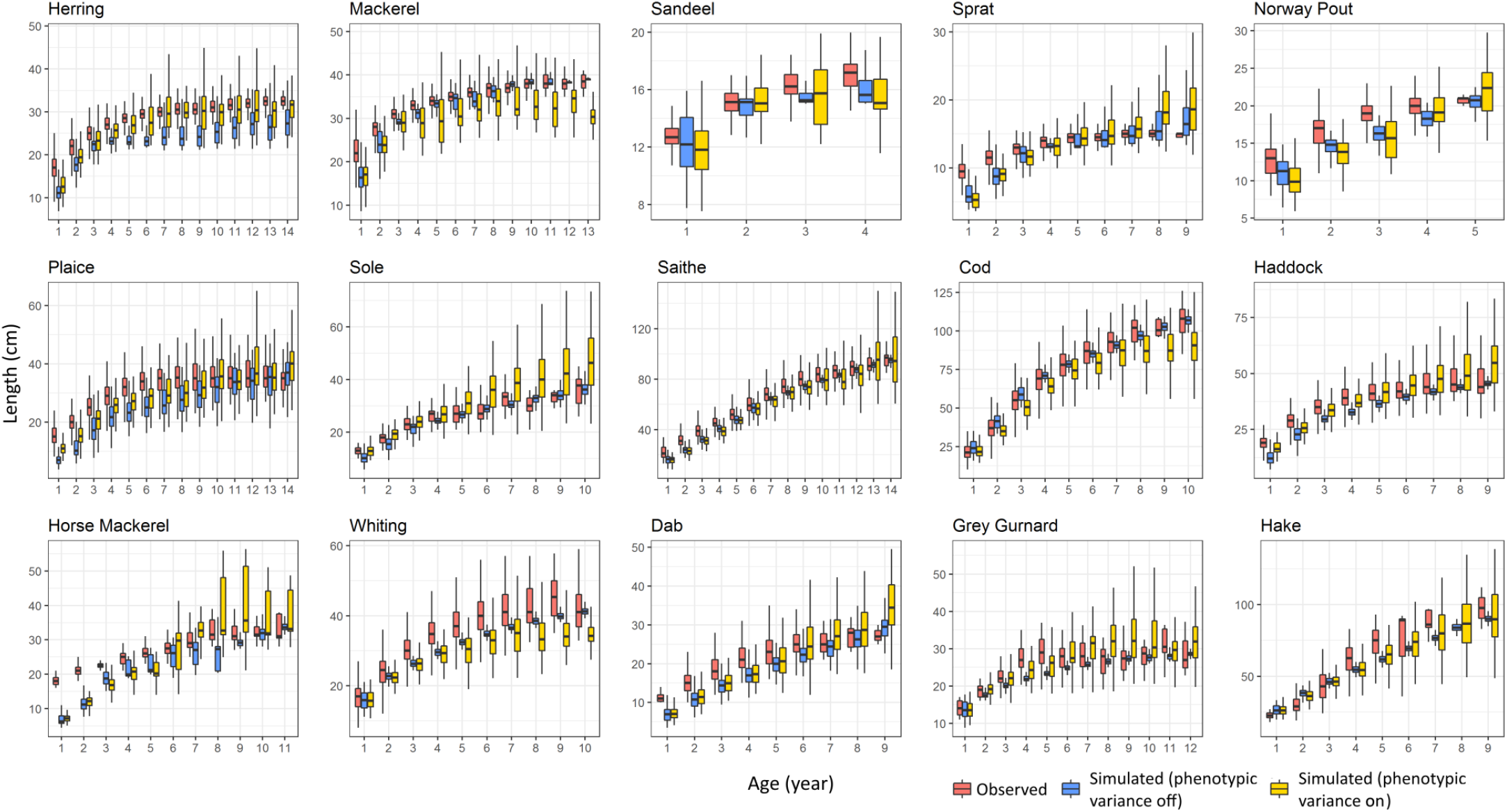
Boxplot of length-at-age per species for observed (red), simulated without (blue) and simulated with phenotypic variance (yellow) individual data. Horizontal bars represent the first, second and third quartiles of the data. The whiskers’ extremities represent 1.5 times the interquartile space (the distance between the first and third quartile).

**Figure 7:**
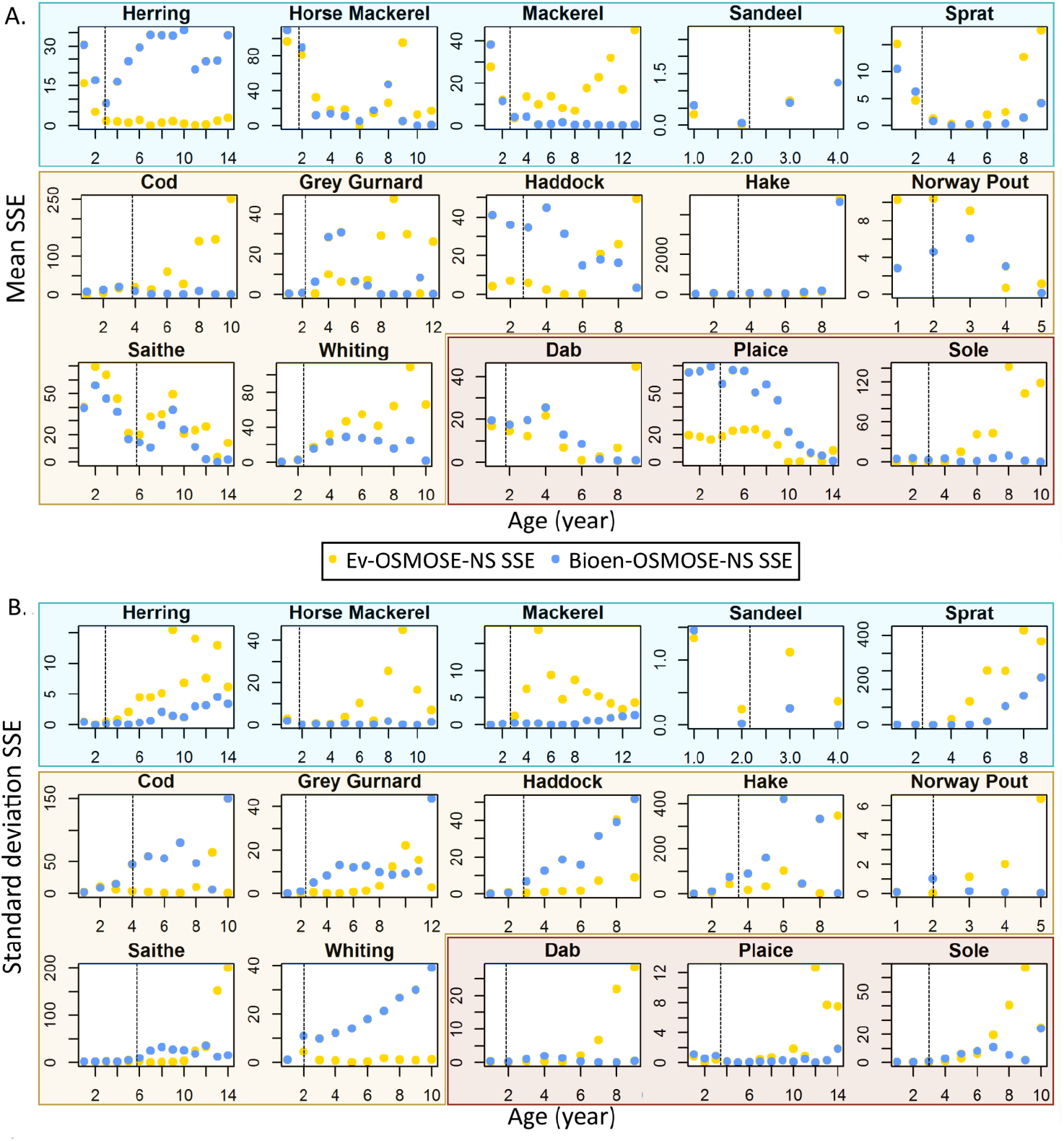
Sum of squared errors between observed and simulated mean (A) and standard deviation (B) of length-at-age from Ev-OSMOSE-NS (yellow dots) and Bioen-OSMOSE-NS (blue dots) per species. The vertical dotted lines represent the mean observed age at maturation. The species are grouped per position in the water column: pelagic (blue frame), demersal (beige frame) and benthic (brown frame) species (see Fig. 4).

The length-at-age outputs from Ev-OSMOSE-NS correctly reproduce the shape of a von Bertalanffy-like growth curve and the length hierarchy between species. Fig. 6 and 7A highlight the degree of similarity in simulations of mean length-at-age between Bioen-OSMOSE-NS and Ev-OSMOSE-NS. Ev-OSMOSE produces better results in terms of mean for herring, haddock, and plaice and fits less well for mackerel, cod, Norway pout, saithe and whiting (Fig. 7A). A recurring trend is that the mean lengths-at-age simulated with Ev-OSMOSE fit poorly observed data for the older ages (cod, dab, grey gurnard, haddock, mackerel, sandeel, sole, sprat, whiting) while the Bioen-OSMOSE-NS results fit better at these ages.

We highlight three main trends in the fit of our models to the observed variability of length-at-age (Fig. 6 and 7B): (i) Ev-OSMOSE outputs generally fit better variance in observed data than Bioen-OSMOSE outputs for demersal species (in particular cod, haddock, saithe, whiting), except Norway pout, but (ii) not for pelagic species (herring, mackerel, sandeel, horse mackerel, sprat) and (iii) the fit is better at earlier ages than at older ages, i.e. the Ev-OSMOSE results tend to overestimate the variance of length at older ages.

### 3.2. Genotypic value transmission

To simulate evolution, a part of the phenotypic variability needs to be transmitted from parents to offspring through mendelian inheritance. Phenotypic variability was described in part 3.1. Hereafter, we present results that validate the mendelian transmission process. The support of transmission of part of the phenotypic variability is the genotype and more precisely mean genotypic values are transmitted from parental pools to their offspring cohorts thanks to mendelian inheritance of alleles. Figure 8 illustrates the model capacity to transmit the parents’ genotypic value to their offspring for the LMRN intercept *m*_0_ (A) and for the gonado-somatic index *r* (B). This figure shows the linear regression between the fecundity-weighted mean parental genotypic value and the newborn genotypic value for each trait. A perfect transmission occurs when the regression slope is equal to 1 and the regression adjustment (R2) is close to 1. Overall, for the two tested traits, we observe a good transmission of genotypic values. The regression slope is positive for all the species for both traits and between 0.5 and 1.2 for all species, except for herring for *r*. The transmission of *m*_0_ is very good for 4 species (mackerel, sandeel, saithe and grey gurnard). The worst cases for *m*_0_ are observed for sole, haddock and dab. The transmission of *r* is very good for 7 species (mackerel, sandeel, Norway pout, saithe, horse mackerel, grey gurnard and hake). The worst cases for *r* are observed for herring, cod, whiting, and dab. Imperfect transmission of genotypic values is probably due to genetic drift generated by the stochasticity in allele sampling, so-called stochastic sampling error, that could emerge from an insufficient diversity of genotypes in the population (i.e., an insufficient number of schools) or an insufficient number of new produced genotypes (i.e., insufficient number of new born schools). The number of newborn schools per reproductive event is a model parameter (Morell et al., 2023) from which depends the total number of schools of a population. A simulation with 10 times more added schools per reproduction event than in the current configuration is presented in Supporting Information C. The simulated transmission patterns in these additional simulations are less noisy and much closer to perfect transmission of genotypic values between parental populations and their offspring.

**Figure 8:**
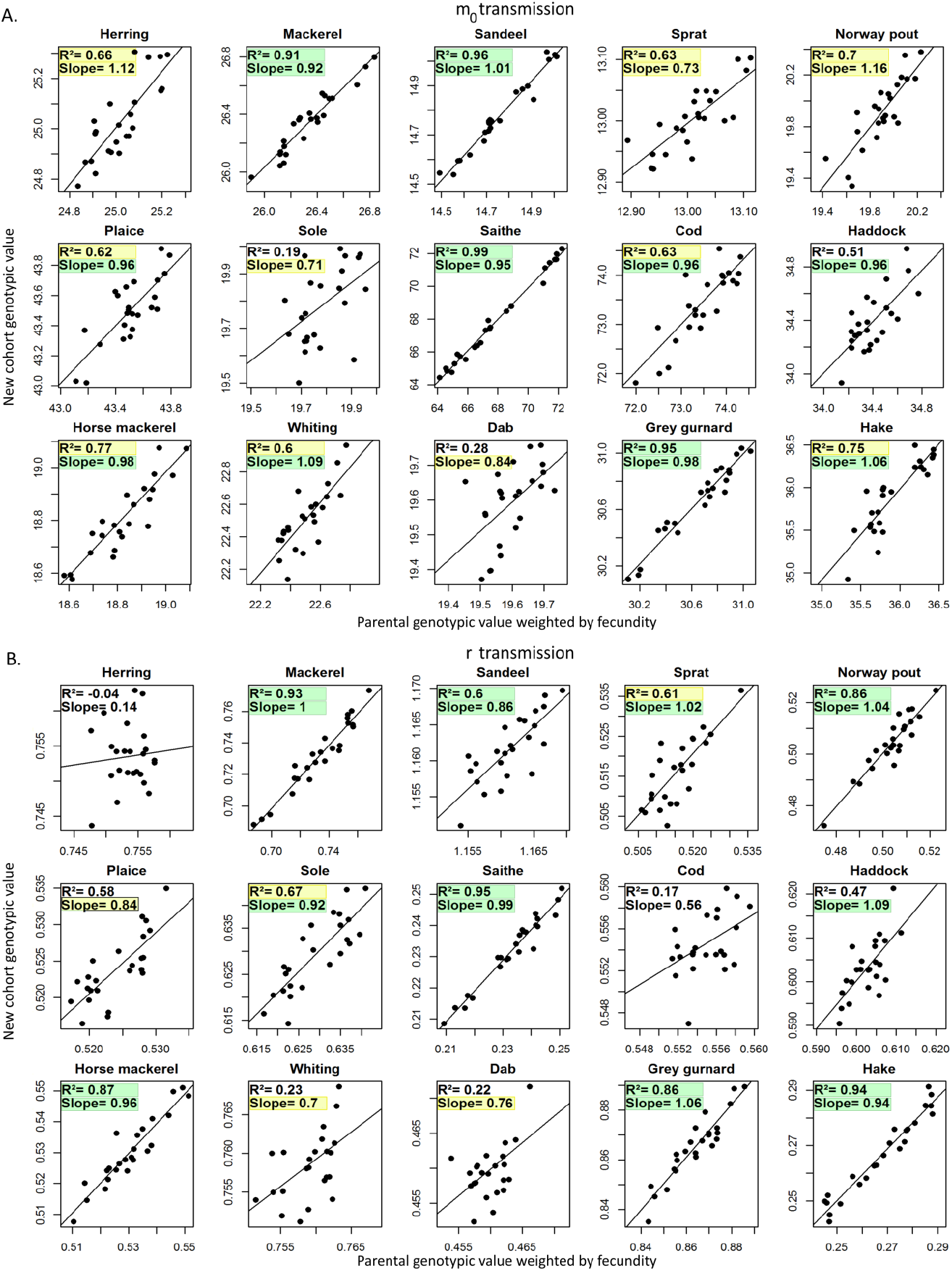
Transmission of genotypic values of the maturation reaction norm intercept *m*_0_ (A) and of the gonado-somatic index *r* (B) from parental pools to the new spawned cohort. The mean parental genotypic value weighted by individual fecundity and averaged over the entire reproductive season is compared to the mean genotypic value of the new spawned cohort during the same reproductive season. The slope and the R^2^ of the regression are expected to be close to 1 in case of faithful transmission of genotypic values. The noise around the regression slope is a consequence of genetic drift due to stochasticity in the sampling of parental alleles. The slope and the R2 highlighted in yellow and green are respectively the faithful and the very good faithful simulated transmission of genotypic values (green slope: between 0.9 and 1.1; yellow slope: between 0.7 and 0.9 or between 1.1 and 1.3; green R2: between 0.8 and 1; yellow R2: between 0.6 and 0.8).

## 4. Discussion

### 4.1. Modeling phenotypic variance of life-history traits

#### 4.1.1. Ev-OSMOSE-NS: A first step to model phenotypic variance

In this study, by applying the evolutionary model Ev-OSMOSE to the North Sea, we obtained a convincing average state of the ecosystem (Supporting Information B, Fig. 5, 6 and 7) and a good overall representation of the variance of life-history traits. The representation of the phenotypic variance is particularly good for the maturation process and encouraging for the growth process (Fig. 5, 6 and 7).

The good representation of the length variance for juveniles to young adults for the majority of species is an indication of a good estimation of the phenotypic variance of the maximum ingestion rate 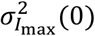 (Fig. 7). Similarly, the good simulated slope of the age- and especially the length-based maturity ogives indicates the reaction norm maturation variance 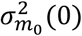 is correctly estimated (Fig. 5). The overestimation of length variance at older ages indicates that (i) one or more aspects impacting these variances still need to be improved in the model, such as assumptions for variance parameter estimates or the reliability of some simulated mechanisms and/or (ii) the quality of length data at older ages is not good enough to be reliable.

#### 4.1.2. Life history parameterization improvement

The mismatch between simulated and observed variance for length at older ages indicates that the simulation of the adult part of life history still needs improvement. The large SSE between simulated and observed adult length-at-age variances is also partly due to the poor data quality at the oldest ages due to a small number of samples. In addition, the data samples are collected on fish that survive until these ages: as fish experience selective pressures over their entire life (mainly fishing), we estimate the input variance of *r* using the surviving fish, i.e., only the surviving genotype/phenotype, which is possibly not representative of the original population diversity required as input.

In other words, the poor data quality implies a poor estimate of the gonado-somatic ratio variance 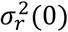 that results in a poor fit between simulated and observed data at older ages, as the observed length-at-age variance is probably lower than it should be. Another source of poor estimation of 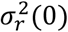 could come from the parameter estimation procedure where we assume that there is no co-variation between *r* and *I_max_*. This hypothesis could be tested using individual growth curves from otolith back-calculation (Green et al., 2009) or data from experimentally raised individuals. Lastly, an incorrect modeling of the foraging-mortality tradeoff would impact the mean length and its variance at adult stage even without evolution: as predation is length-dependent, if the foraging-mortality trade-off does not counterbalance realistically the benefits to grow faster and toward higher lengths, then the simulated phenotypes with a higher *I_max_* survive better and are more abundant at older age than in the wild, overestimating the mean and variance of length, as emerging in simulation from Ev-OSMOSE-NS (Fig. 6, 7).

#### 4.1.3. Prey, predators and fishing impact emerging individual properties

Length-at-age depends on growth, maturation and reproductive parameters as well as size selective pressures such as fishing or predation. For example, an incorrect parameterization of fishing selectivity and a higher simulated exploitation rate than the actual one can lead to a smaller simulated than observed length at adult stage, a pattern that can become even more apparent in Ev-OSMOSE-NS when more phenotypic variability is added in the population. This case is observed for mackerel, sandeel and cod for example (Fig. 6). The simulated lengths-at-age show a decrease at older ages. This pattern emerges from the truncation of the fast-growing fish part of the population: the fish that survive to these ages are small and slow-growing. If this pattern is not observed in the data, it reflects overfishing in the simulation, either in terms of total fishing pressure or selectivity for larger lengths. The addition of growth process variability accentuates this pattern.

#### 4.1.4. Limits from the model’s life history description

The observed length-at-age variance is the sum of the variances due to additive genetic variability, the phenotypic expression noise and the phenotypic plasticity emerging from macro-environmental variations (Fig. 2). In our method to estimate process-based-trait variance, we assumed that the emerging variance was the result of additive genetic and phenotypic expression noise variances only. Thus, the model performs better on species for which phenotypic plasticity in response to macro-environmental variations has few impacts on length-at-age variance such as cod, whiting, saithe or haddock for example (blue boxplots in Fig. 6 and variance SSE in Fig. 7). On the contrary, this implies that the simulated length-at-age variance is overestimated for species with a high phenotypic plasticity variance emerging from macro-environment variations in the wild. These species are mainly the small pelagic species (herring, sprat, and sandeel mainly) that feed on highly variable sources of food, mainly phyto- and zooplankton. Accounting for macro-environmental variations in variance parameter estimations would be a way to improve the simulated length-at-age variance.

The assumption of linearity for the maturation reaction norm does not allow to correctly represent the maturation patterns for some species such as mackerel (Fig. 5A). By contrast to other species, the slope of the LMRN of mackerel is positive: fish that mature older are bigger. The species that empirically exhibit this maturation pattern also frequently exhibit a reaction norm that decreases at older ages (Heino et al., 2002; Marty et al., 2014) or a maximum length to mature (Nilsson-Örtman and Rowe, 2021). With a strictly increasing LMRN, some individuals never mature if their LMRN slope is steeper than their growth rate. This case was not observed in Bioen-OSMOSE-NS (i.e. without phenotypic variance) but appears here in Ev-OSMOSE-NS with the modeling of phenotypic variance generating some individuals combining a steep positively sloped LMRN and slow growth.

#### 4.1.5. Toward more evolving traits: technical improvement

In this study, we presented the simulated effects of phenotypic variance on three process-based traits, with activation of evolution on two of these traits, i.e., the reproductive investment trait *r* and a maturation process trait *m*_0_. Reproductive investment evolution (Wright et al., 2011; Yoneda and Wright, 2004) and maturation evolution (de Roos et al., 2006; Marty et al., 2014; Mollet et al., 2007) are the two main known consequences of fisheries-induced evolution. Reported length-at-age evolution in the literature (Enberg et al., 2012) can be the consequence of evolutionary changes in the reproductive investment, the maturation process or juvenile growth. The evolution of juvenile growth was not modeled here in agreement with the fact that it has been seldom documented and remains weak compared to other traits’ evolution (Enberg et al., 2012; Heino et al., 2015). Moreover, to correctly model juvenile growth evolution, which in our model translates into maximum mass-specific ingestion rate *I_max_* evolution, the account of a trade-off between the foraging intensity, that should be positively related to *I_max_*, and its associated mortality *M*_f_ is necessary (Enberg et al., 2009) but is difficult to parameterize in the absence of in situ or experimental data. A way to parameterize this trade-off in future studies would be to estimate the *M*_f_ unknown parameters *k*_1_ and *k*_2_ by using time series of trait values in an hindcast interannual calibration. Including the evolution of *I_max_* would greatly increase the realism of the model as evolutionary pressures impact multiple traits including growth, especially in the context of length-selective fishing.

### 4.2. Genotypic value transmission

#### 4.2.1. A good transmission of genotypic values implies a correct evolutionary trend

The genotypic value transmission between parental populations and new cohorts is essential in any eco-evolutionary model such as Ev-OSMOSE because it ensures that the advantageous alleles will be transmitted from parents to offspring: the effect of selection can then propagate through generations.

The transmission is validated from Figure 8 and Supplementary Information C, as we observed that the fecundity-weighted mean genotypic values of the parental pools are transferred to the newborn cohort. Furthermore, at the species level, a larger number of schools improves genotypic value transmission (see Supplementary Information C), decreasing the noise by reducing alleles’ stochastic sampling error and thus genetic drift (see 4.2.2).

Obtaining positive slopes and high R^2^ for regressions of newborn genotypic values on parental ones indicates faithful transmission for both traits and for all the species. Then, the resulting evolutionary trends are reliable in terms of response to selection: a change in a parental trait’s genotypic value due to selection during parent lifetime will be transmitted to offspring. The difference between a species with a faithful transmission and a species with a noisy one, as long as the slope is positive, will be in terms of the rate of the evolutionary response: the stochasticity in transmission will slow down the evolutionary response.

#### 4.2.2. Genetic drift: a model sensitive to the number of super-individuals (schools)?

In the Ev-OSMOSE model, and more generally in the OSMOSE model, the biological individuals (fish) are grouped in super-individuals (groups of fish, called schools) to improve the calculation time. In the model, the number of schools added per reproductive event is empirically fixed to have at minimum a school of each age class per species per cell where the species is distributed. This minimum number of schools is a trade-off between reducing the stochasticity of the model and decreasing the computing needs (both in terms of required memory and calculation time). The use of a genetic sub-model that explicitly describes the genetic diversity in the population implies another condition to determine the minimum number of schools, which is to limit stochasticity in allele sampling during reproductive events and then genetic drift. The genetic drift is related to the population size (in our case the number of schools per species) as it decreases with it (Masel, 2011) and more precisely with the associated effective population, defined as the size of an ideal population (random mating, equal sex ratio and no overlapping generations) that would have the same rate of genetic change than the actual population (Beissinger and McCullough, 2002). In Ev-OSMOSE, the structure in school limits the maximum effective population size at the number of schools and not the total abundance of individuals, which could artificially increase genetic drift.

These considerations are highlighted by comparing a simulation with a lower number of schools (Fig. 8) that displays a stronger genetic drift than a simulation with 10 times more schools added per reproductive event (Supporting information C). The increase of the number of schools in Ev-OSMOSE is limited due to problems in terms of calculation time: 50 years of the configuration presented in this paper runs in 20 minutes whereas 50 years of the configuration presented in Supporting Information C where the only difference is the number of schools runs in 15 hours on the same computer. Knowing that the model needs to be run thousands of times to be calibrated, this difference in calculation time cannot be neglected. It would be necessary to conduct a sensitivity analysis to identify an acceptable compromise between the faithfulness of genotypic value transmission, genetic drift and calculation time.

An interesting aspect is also the difference of genotypic value transmission between species. Some species exhibit an almost perfect transmission with a low number of schools (e.g., saithe, Fig. 8) whereas others are still very noisy in the simulation with a high number of schools (e.g. whiting, Supporting Information C). We hypothesize that differences at the interspecific level could arise from differences between species in terms of demography, selective pressures or genetic structure. Regarding the demography, the total size of the population, the total number of schools in the population, the number of schools added per reproductive event and the total fecundity were not correlated with the faithfulness of the transmission (results not shown). The age structure of the mature part of the population could be an interesting feature to explore as overlapping reproductive generations partly explains differences between effective and real population sizes, and is the only source of differences between these included in Ev-OSMOSE, as otherwise mating is random and sex ratio is balanced. Regarding the genetic structure of the population, we observed than genotypic and phenotypic variances, heritability, allele frequencies and heterozygosity were not correlated with the faithfulness of transmission (results not shown). A next step would be to explore the relationship with effective population size and genetic grift. Lastly, as genetic drift impact is expected to be stronger for small populations or weak selection (Barton and Partridge, 2000), it would be interesting to explore the link between selective pressure intensity and genetic drift.

## 5. Conclusion

This first application of the eco-evolutionary multi-species model Ev-OSMOSE to the North Sea opens the field of eco-evolutionary studies to marine ecosystems models. This study underlines the parameterization feasibility in spite of the high data quality requirement to parameterize the phenotypic and genotypic variances of life-history traits. Ev-OSMOSE-NS is the first configuration to account for genotypic and phenotypic variances of several interacting species and succeeds to improve the simulated variances of life-history traits. It is an important step toward more realism notably in representing length-at-age distribution and the maturation process.

Ev-OSMOSE-NS is also, to our knowledge, the first multi-species model applied to a marine ecosystem that accounts for mendelian inheritance of traits from parents to their offspring for all the species of a food web simultaneously, thus allowing to account for the micro-evolution of exploited species in response to selective pressures such as fishing and climate change together with their co-evolution due to trophic interactions.

A next step is to use the Ev-OSMOSE model under climate change or fishing scenarios. We believe that the account of eco-evolutionary dynamics will improve future projections of marine biodiversity, at the interspecific and intraspecific levels, and fulfill a gap of knowledge on the evolution of interacting species in communities under multiple natural and anthropogenic selective pressures.

## Acknowledgements

This work has been partially funded by the BiodivErsA and Belmont Forum project SOMBEE (BiodivScen programme, ANR contract n°ANR-18-EBI4-0003-01). Alaia Morell was supported by a PhD grant from Ifremer and Région Hauts-de-France. The POLCOMS-ERSEM projections were produced with funding from the European Union Horizon 2020 research and innovation programme under grant agreement No 678193 (CERES, Climate Change and European Aquatic Resources). The authors acknowledge the Pôle de Calcul et de Données Marines (PCDM, http://www.ifremer.fr/pcdm) for providing DATARMOR storage, data access, computational resources, visualization and support services. Yunne-Jai Shin acknowledges funding support from the European Union’s Horizon 2020 research and innovation program under grant agreement No 869300 (FutureMARES) and the Pew marine fellows program.

## Author Contributions

Yunne-Jai Shin and Bruno Ernande conceived and supervised the project. Bruno Ernande and Alaia Morell conceived the concepts of the new model developments. Nicolas Barrier and Alaia Morell developed the code and validated the model functioning. Alaia Morell gathered the data for the model parameterization. Bruno Ernande and Alaia Morell conceived and developed the scripts for the parameter estimation. Morgane Travers and Alaia Morell parameterized the model. All authors interpreted the model outputs. Nicolas Barrier, Morgane Travers and Alaia Morell performed the model calibration. Alaia Morell wrote the first paper draft. All authors contributed critically to the revisions of the manuscript and gave final approval for submission.

## Data Archiving

The Ev-OSMOSE-NS configuration and its associated version of the Ev-OSMOSE model executable will be deposited on Zenodo. Model code will be available on Github. The scripts developed to estimate Bioen-OSMOSE-NS parameters are available on Github.

## 7. Supporting Information

### Supporting Information A Estimation procedure for the coefficient of variations of the traits under selection

#### A.1. Estimation of phenotypic variance of the juvenile growth coefficient *c* and the gonado-somatic index *r*

The growth in length of an individual *i* can be described as

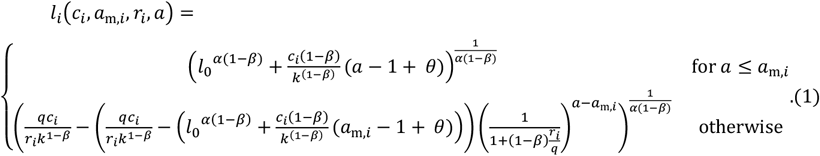

We denote 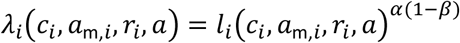 the transformed length at age, which gives

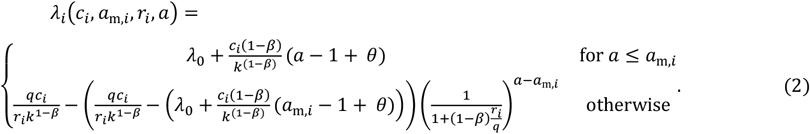

with 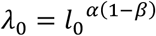.

Applying the Delta method to the growth equation (2) and neglecting second order terms in the Taylor expansion, we obtain

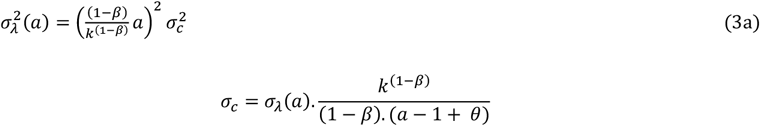

for any age *a* ≤ *a*_m_ and

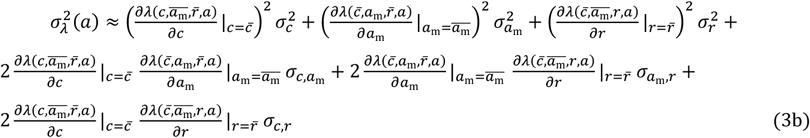

for any age *a* > *a*_m_ where 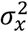 and *σ_x, y_* denote variance of *x* and covariance of *x* and *y*, respectively Under the assumption of negligible covariances between *c, a_m_*, and *r* we obtain further

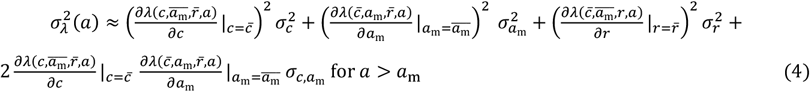

Denoting the age-dependent maturity ogive *o*(*a*), the maturation probability at age is obtained as

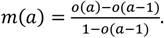

Mean maturation can thus be obtained as

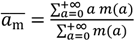

and its variance as

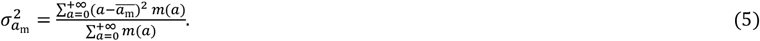

Given that we know 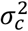 from equation (3a) and 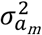 from equation (5), then we can deduce an approximation of 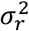 from equation (4)

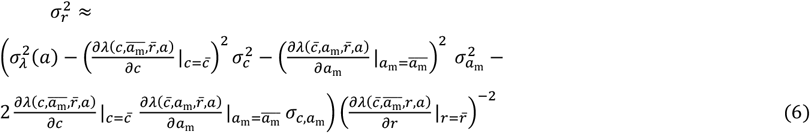

Equations (3a) and (6) allow estimating 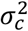 and 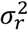 from length-at-age data for fully immature individuals, i.e., for any age *a* so that *o*(*a*) = 0 for the former, and for fully mature individuals, i.e., for any age *a* so that *o*(*a*) = 1 for the latter.

One way to combine these equations for an estimation across all ages is to look for the values of 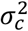 and 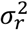 minimizing the sum of squared differences between 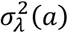, the variance of transformed length at age, and the right handside of equations (3a) and (4) respectively weighted by the probability of being immature (1 – *o*(*a*)) and being mature *o*(*a*):

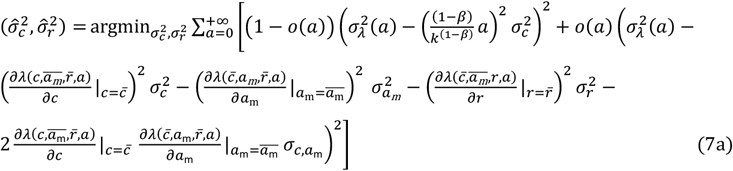

with 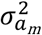 estimated using equation (5) and with

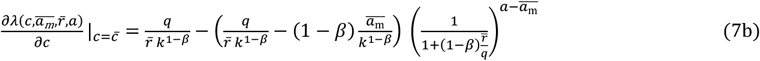

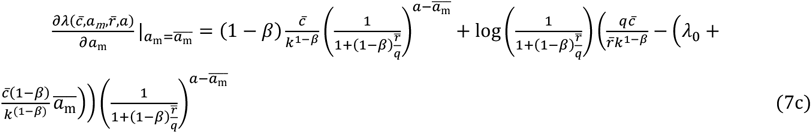

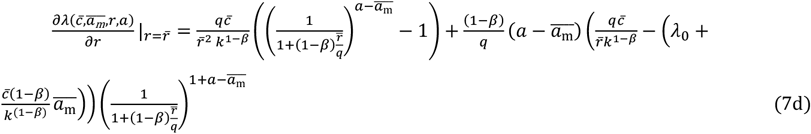

where 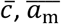, and 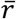 are the mean parameter values that can were estimated from the input parameter estimation procedure.

#### A.2 Estimation of phenotypic variance of the linear probabilistic maturation reaction norm parameters

The maturation probability of an individual *i* of age *a* and length *l* conditional on being alive and still immature can be described by a Heaviside step function

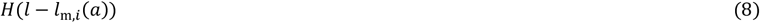

where *l*_m,*i*_(*a*) is the individual’s maturation length at age *a*. Equation (8) thus describes an individual’s maturation reaction norm. Phenotypic variation in maturation length across individuals aged *a* is described by the probability density function *f_a_*(*l*_m_) with mean 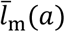 and standard deviation *σ*_*l*_m__(*a*). The population-level PMRN *p*(*l, a*) is then obtained as

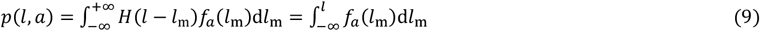

which is the cumulative distribution function of maturation lengths at age *a*.

The derivative of the population-level PMRN according to length allows thus to empirically estimate the probability density function of maturation length

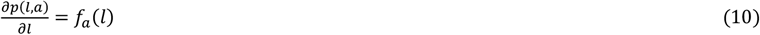

The mean and variance of maturation length at any age *a* can thus be estimated from the empirical PMRN 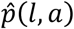 as

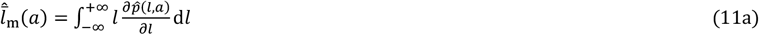

and

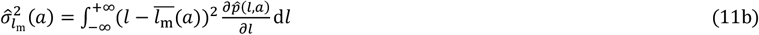

Under the assumption of a linear maturation reaction norm with a fix envelop, the maturation length of an individual *i* is described at any age *a* by

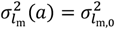

#### A.3. Estimation of covariance between the juvenile growth coefficient c and age at maturation a_m_

To compute the covariance between age a maturation *a*_m_ and growth potential *c*, we need to estimate the joint probability density of these two random variables that we will denote as *g*(*a*_m_, *c*).

Under the assumptions of our model, i.e. that survival only depends on length, the number of newly maturing individuals between age *a* and *a* + 1 and between transformed length *λ* = *l*^*α*(1-*β*)^ and *λ* + Δ*λ* is given by

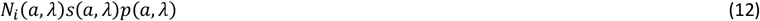

where *N_i_*(*a,λ*) is the number of immature individuals aged *a* with transformed length *λ*, *s*(*a, λ*) is survival from age *a* to *a* + 1 and transformed length *λ* to *λ* + Δ*λ* (which results from the combination of natural and fishing mortality) and *p*(*a, λ*) is the prospective version of the PMRN.

As immature growth in transformed length *λ* is linear with age according to 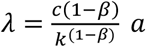, all dependencies on *λ* can be turned into dependencies on *c* by a simple change of variable

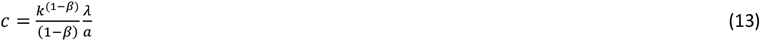

so that the number of newly maturing individuals between age *a* and *a* + 1 for a growth potential *c* is given by

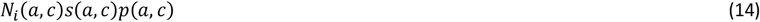

If the age and length distribution of sampled individuals *n*_i_ is representative of that of the population *N*_i_, the joint probability distribution of maturation age and growth potential is then obtained as

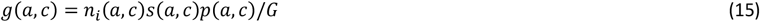

with 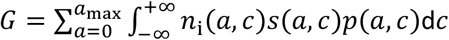 a normalization constant insuring that the joint probability density function sums to 1.

An estimate of the covariance between age at maturation *a*_m_ and growth potential *c* is then obtained as

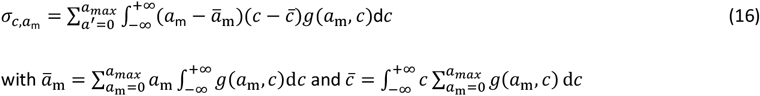

### Supporting Information B Ecological validation of Ev-OSMOSE-NS: Simulated biomass and catches

**Figure S1:**
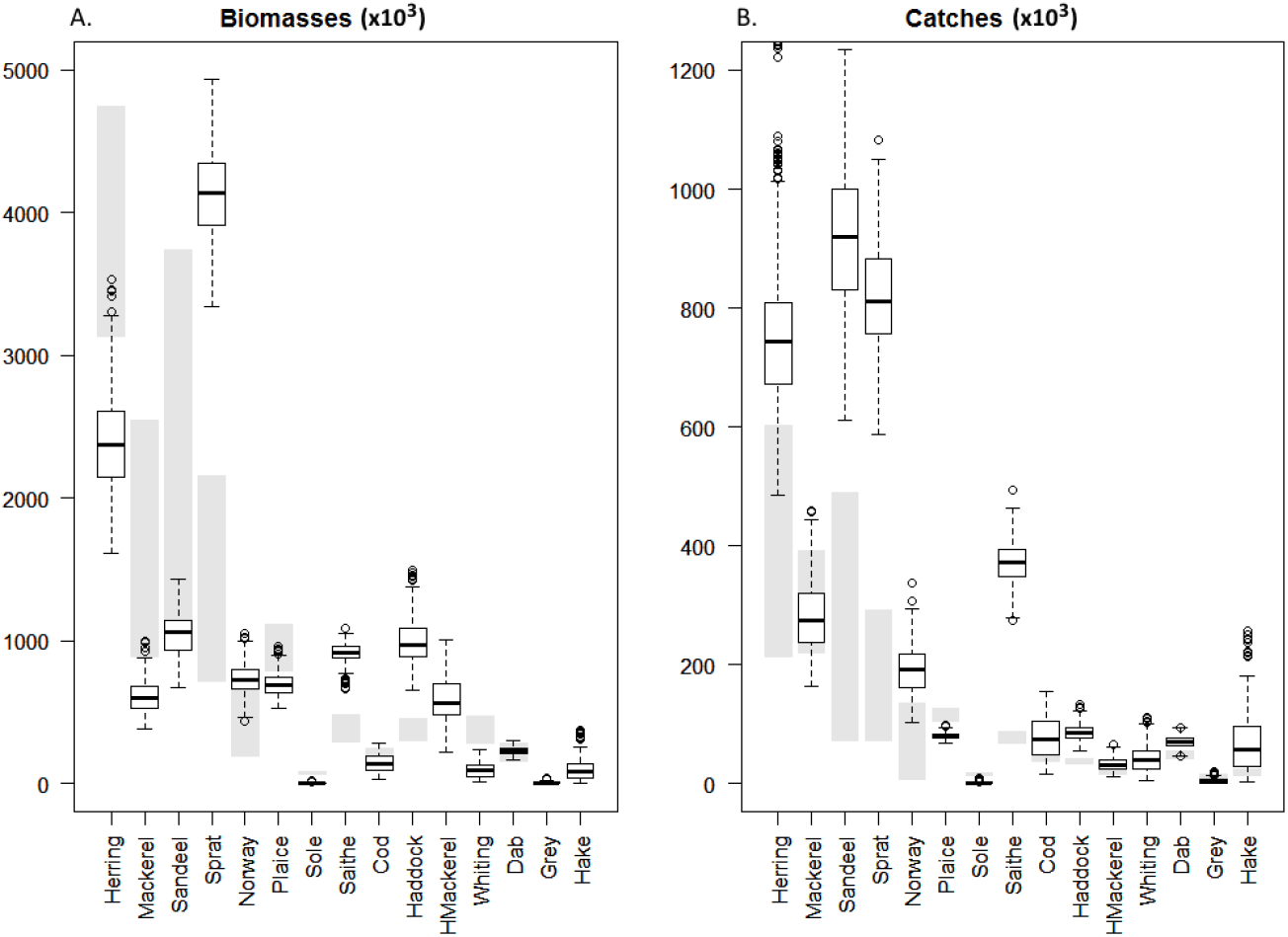
Fisheries catches (A) and biomasses (B), in thousand tons, per species for stock assessment estimates and simulated data averaged over 28 replicates (boxplots). The boxplots represent the simulated data for 28 replicated simulations (stochastic model) for the catches and biomasses per species, with the first, second and third quartiles represented horizontally in each plot. The averaged simulated are from year 50 to 70, before the evolution activated at the year 70. The gray bars show the minimal and maximum values observed for catch and biomass estimates from stock assessment for the 2010-2019 period. The species without gray bars for biomasses are not assessed in the area.

**Figure S2:**
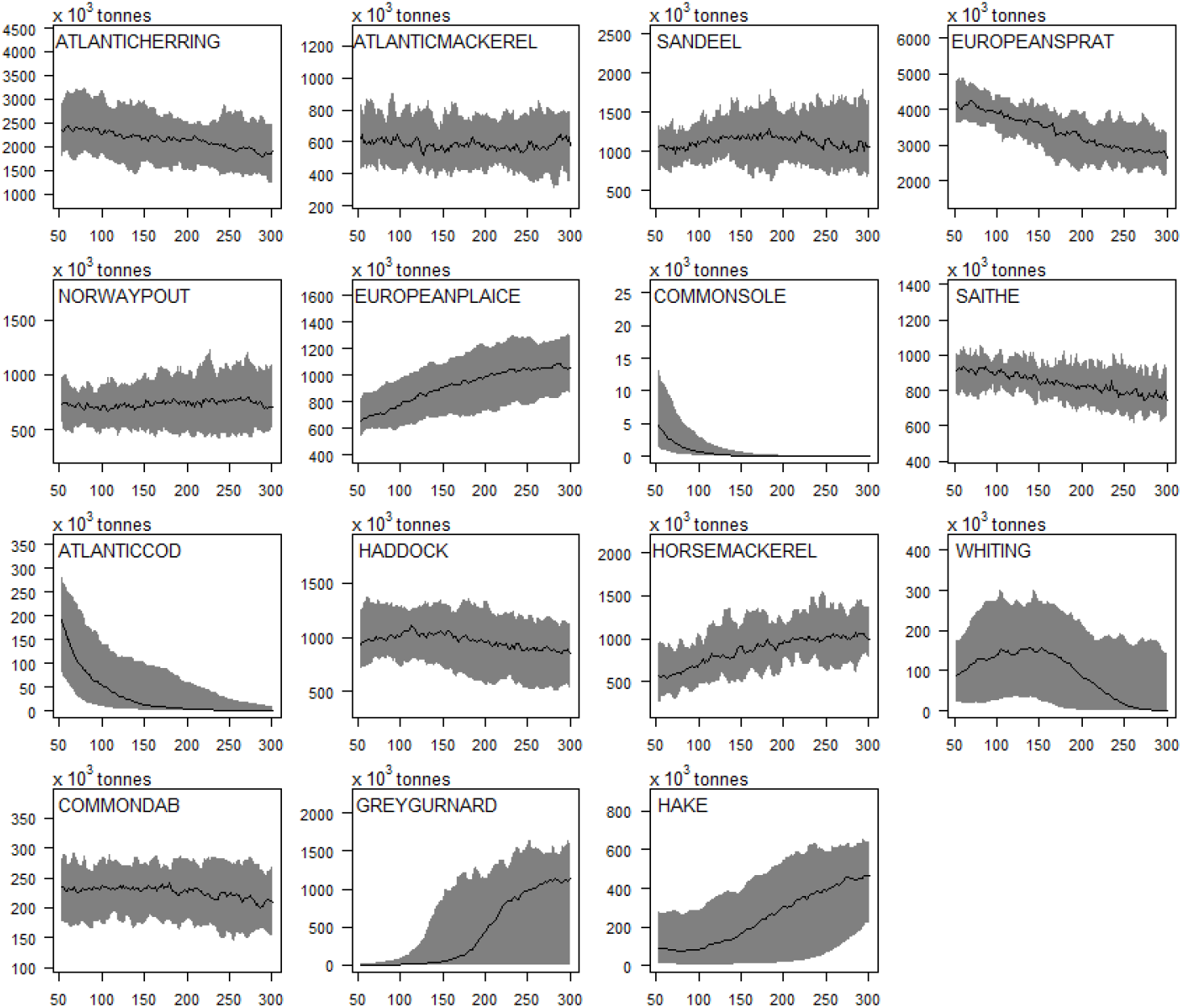
Simulated time series of biomasses. Data averaged over 28 replicates (black line) and replicates variability due to stochasticity (grey area). The configuration is considered stable between year 50 and 70, except for cod and sole. The genotype transmission is activated after year 70.

### Supporting Information C Genotype transmission validation

**Figure S2:**
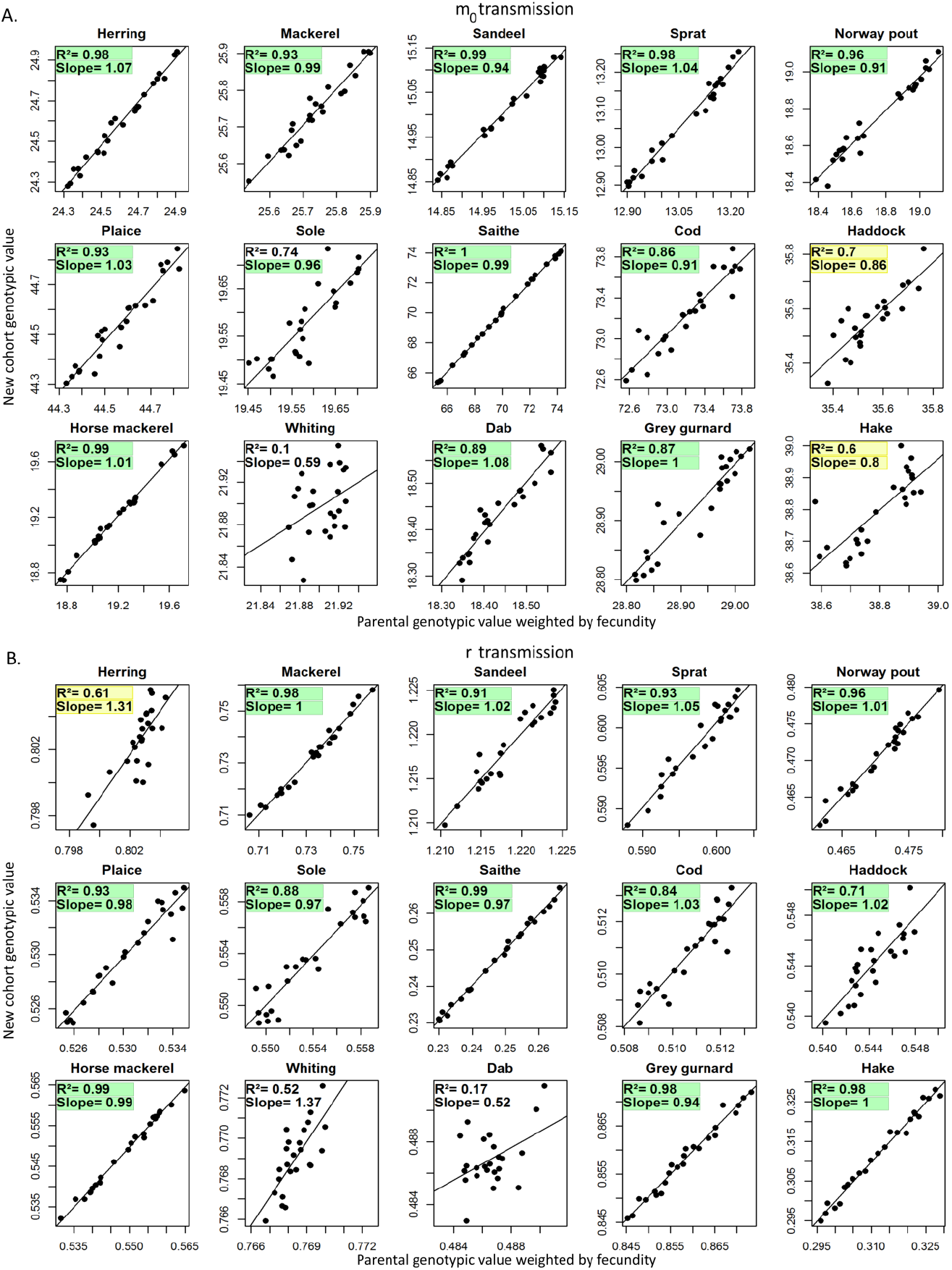
Transmission of genotypic value of the maturation reaction norm origin m0 (A) and of the gonado-somatic index r (B) from parent to the new spawned cohort in a simulation where the number of schools created per reproductive event is 10 times higher than in the configuration presented in the main text. The mean parent genotypic value weighted by individual fecundity average over the entire reproductive season time step is compared to the mean genotypic value of the new spawned cohort during the same reproductive season. The slope and the regression adjustment are expected to be close to 1. The noise around the regression slope is a consequence of drift due random parental allele selection and random mating. As in figure 8, The slope and the R^2^ highlighted in yellow and green are respectively the good and the very good fit of simulated data to expected pattern (green slope: between 0.9 and 1.1; yellow slope: between 0.7 and 0.9 or between 1.1 and 1.3; green R2: between 0.8 and 1; yellow R2: between 0.6 and 0.8).

## References

Andersen, K.H., 2019. Fish Ecology, Evolution, and Exploitation: A New Theoretical Synthesis. Princeton University Press.

Audzijonyte, A., Kuparinen, A., Fulton, E., 2014. Ecosystem effects of contemporary life-history changes are comparable to those of fishing. Marine Ecology Progress Series 495, 219–231. https://doi.org/10.3354/meps10579

Audzijonyte, A., Kuparinen, A., Gorton, R., Fulton, E.A., 2013. Ecological consequences of body size decline in harvested fish species: positive feedback loops in trophic interactions amplify human impact. Biology Letters 9, 20121103–20121103. https://doi.org/10.1098/rsbl.2012.1103

Barton, N., Partridge, L., 2000. Limits to natural selection. Bioessays 22, 1075–1084. https://doi.org/10.1002/1521-1878(200012)22:12<1075::AID-BIES5>3.0.CO;2-M

Beissinger, S.R., McCullough, D.R., 2002. Population Viability Analysis. University of Chicago Press.

Boukal, D.S., Dieckmann, U., Enberg, K., Heino, M., Jørgensen, C., 2014. Life-history implications of the allometric scaling of growth. Journal of Theoretical Biology 359, 199–207. https://doi.org/10.1016/j.jtbi.2014.05.022

Crozier, L.G., Hutchings, J.A., 2014. Plastic and evolutionary responses to climate change in fish. Evolutionary Applications 7, 68–87. https://doi.org/10.1111/eva.12135

de Roos, A.M., Boukal, D.S., Persson, L., 2006. Evolutionary regime shifts in age and size at maturation of exploited fish stocks. Proceedings of the Royal Society B: Biological Sciences 273, 1873–1880. https://doi.org/10.1098/rspb.2006.3518

Dunlop, E.S., Heino, M., Dieckmann, U., 2009. Eco-genetic modeling of contemporary life-history evolution. Ecological Applications 19, 1815–1834. https://doi.org/10.1890/08-1404.1

Enberg, K., Jørgensen, C., Dunlop, E.S., Heino, M., Dieckmann, U., 2009. ORIGINAL ARTICLE: Implications of fisheries-induced evolution for stock rebuilding and recovery: Fisheries-induced evolution and stock recovery. Evolutionary Applications 2, 394–414. https://doi.org/10.1111/j.1752-4571.2009.00077.x

Enberg, K., Jørgensen, C., Dunlop, E.S., Varpe, Ø., Boukal, D.S., Baulier, L., Eliassen, S., Heino, M., 2012. Fishing-induced evolution of growth: concepts, mechanisms and the empirical evidence: Fishing-induced evolution of growth. Marine Ecology 33, 1–25. https://doi.org/10.1111/j.1439-0485.2011.00460.x

Green, B.S., Mapstone, B.D., Carlos, G., Begg, G.A., 2009. Tropical Fish Otoliths: Assessment, Management, and Ecology.

Heino, M., Díaz Pauli, B., Dieckmann, U., 2015. Fisheries-Induced Evolution. Annual Review of Ecology, Evolution, and Systematics 46, 461–480. https://doi.org/10.1146/annurev-ecolsys-112414-054339

Heino, M., Dieckmann, U., Godø, O.R., 2002. Measuring probabilistic reaction norms for age and size at maturation. Evolution 56, 669–678.

Heymans, J.J., Bundy, A., Christensen, V., Coll, M., de Mutsert, K., Fulton, E.A., Piroddi, C., Shin, Y.-J., Steenbeek, J., Travers-Trolet, M., 2020. The Ocean Decade: A True Ecosystem Modeling Challenge. Frontiers in Marine Science 7. https://doi.org/10.3389/fmars.2020.554573

Lynch, M., Walsh, B., 1998. Genetics and Analysis of Quantitative Traits, Sinauer Associates, Inc, Sunderland. ed.

Mangel, M., 2003. Environment and Longevity: The Demography of the Growth Rate 15.

Marty, L., Rochet, M., Ernande, B., 2014. Temporal trends in age and size at maturation of four North Sea gadid species: cod, haddock, whiting and Norway pout. Marine Ecology Progress Series 497, 179–197. https://doi.org/10.3354/meps10580

Masel, J., 2011. Genetic drift. Current Biology 21, R837–R838. https://doi.org/10.1016/j.cub.2011.08.007

Mollet, F.M., Kraak, S.B.M., Rijnsdorp, A.D., 2007. Fisheries-induced evolutionary changes in maturation reaction norms in North Sea sole Solea solea. Marine Ecology Progress Series 351, 189–199. https://doi.org/10.3354/meps07138

Morell, A., Shin, Y.-J., Barrier, N., Travers-Trolet, M., Halouani, G., Ernande, B., 2023. Bioen-OSMOSE: A bioenergetic marine ecosystem model with physiological response to temperature and oxygen. https://doi.org/10.1101/2023.01.13.523601

Mousseau, T., Roff, D., 1987. Mousseau TA, Roff DA. Natural selection and the heritability of fitness components. Heredity 59: 181-197. Heredity 59 (Pt 2), 181–97. https://doi.org/10.1038/hdy.1987.113

Naish, K.A., Hard, J.J., 2008. Bridging the gap between the genotype and the phenotype: linking genetic variation, selection and adaptation in fishes. Fish and Fisheries 9, 396–422. https://doi.org/10.1111/j.1467-2979.2008.00302.x

Nilsson-Örtman, V., Rowe, L., 2021. The evolution of developmental thresholds and reaction norms for age and size at maturity. PNAS 118. https://doi.org/10.1073/pnas.2017185118

Poulsen, N.A., Nielsen, E.E., Schierup, M.H., Loeschcke, V., Grønkjær, P., 2006. Long-term stability and effective population size in North Sea and Baltic Sea cod (Gadus morhua). Molecular Ecology 15, 321–331. https://doi.org/10.1111/j.1365-294X.2005.02777.x

Quince, C., Abrams, P.A., Shuter, B.J., Lester, N.P., 2008. Biphasic growth in fish I: Theoretical foundations. Journal of Theoretical Biology 254, 197–206. https://doi.org/10.1016/j.jtbi.2008.05.029

Roff, D.A., 1992. The evolution of life histories: theory and analysis. Chapman and Hall, New York.

Rose, K.A., Allen, J.I., Artioli, Y., Barange, M., Blackford, J., Carlotti, F., Cropp, R., Daewel, U., Edwards, K., Flynn, K., Hill, S.L., HilleRisLambers, R., Huse, G., Mackinson, S., Megrey, B., Moll, A., Rivkin, R., Salihoglu, B., Schrum, C., Shannon, L., Shin, Y.-J., Smith, S.L., Smith, C., Solidoro, C., St. John, M., Zhou, M., 2010. End-To-End Models for the Analysis of Marine Ecosystems: Challenges, Issues, and Next Steps. Marine and Coastal Fisheries 2, 115–130. https://doi.org/10.1577/C09-059.1

Shin, Y.-J., Shannon, L., Cury, P., 2004. Simulations of fishing effect on the southern benguela fish community using an individual-based model : learning from a comparison with ecosim. African Journal of Marine Science 26, 95–114.

Soularue, J.P., Kremer, A., 2012. Assortative mating and gene flow generate clinal phenological variation in trees. BMC Evolutionary Biology, 12, 79.

Stearns, S.C., 1992. The Evolution of Life Histories. OUP Oxford.

Stearns, S.C., Koella, J.C., 1986. The Evolution of Phenotypic Plasticity in Life-History Traits: Predictions of Reaction Norms for Age and Size at Maturity. Evolution 40, 893–913. https://doi.org/10.2307/2408752

Waples, R.S., Audzijonyte, A., 2016. Fishery-induced evolution provides insights into adaptive responses of marine species to climate change. Frontiers in Ecology and the Environment 14, 217–224. https://doi.org/10.1002/fee.1264

Wright, P.J., Gibb, F.M., Gibb, I.M., Millar, C.P., 2011. Reproductive investment in the North Sea haddock: temporal and spatial variation. Marine Ecology Progress Series 432, 149–160. https://doi.org/10.3354/meps09168

Yoneda, M., Wright, P.J., 2004. Temporal and spatial variation in reproductive investment of Atlantic cod Gadus morhua in the northern North Sea and Scottish west coast. Mar. Ecol. Prog. Ser. 276:237–248.

